# Time of day and genotype sensitivity adjust molecular responses to temperature stress in sorghum

**DOI:** 10.1101/2023.04.09.536181

**Authors:** Titouan Bonnot, Impa Somayanda, S. V. Krishna Jagadish, Dawn H Nagel

## Abstract

Sorghum is one of the four major C4 crops that are considered to be tolerant to environmental extremes. Sorghum shows distinct growth responses to temperature stress depending on the sensitivity of the genetic background. About half of the transcripts in sorghum exhibit diurnal rhythmic expressions emphasizing significant coordination with the environment. However, an understanding of how molecular dynamics contribute to genotype-specific stress responses in the context of the time of day is not known. We examined whether temperature stress and the time of day impact the gene expression dynamics in cold-sensitive and tolerant and heat-sensitive and tolerant sorghum genotypes. We found that time of day is highly influencing the temperature stress responses, which can be explained by the rhythmic expression of most thermo-responsive genes. This effect is more pronounced in thermo-tolerant genotypes, suggesting a stronger regulation of gene expression by the time of day and/or by the circadian clock. Genotypic differences were mostly observed on average gene expression levels, but we identified groups of genes regulated by temperature stress in a time-of-day and genotype-specific manner. These include transcriptional regulators and several members of the Ca^2+^-binding EF-hand protein family. We hypothesize that expression variation of these genes between genotypes may be responsible for contrasting sensitivities to temperature stress in tolerant vs susceptible sorghum varieties. These findings offer a new opportunity to selectively target specific genes in efforts to develop climate-resilient crops based on their time of day and genotype variation responses to temperature stress.

## Introduction

To better attune with their environment, plants partition specific responses to the most optimal times of the day. This regulation involves the coordination between the external environment, the circadian clock, internal cellular processes, and biological outputs (Nagel & Kay 2012; McClung 2019). In plants, the circadian clock controls a large portion of the transcriptome that is responsive to stress stimuli (Covington, Maloof, Straume, Kay & Harmer 2008; Markham & Greenham 2021). A subset of this stress-responsive transcriptome is subjected to gating or modulation of gene expression meaning that the transcript abundance of genes varies not only in response to the stress but depending on the time of day it is perceived (Hotta *et al*. 2007; Grundy, Stoker & Carré 2015; Paajanen, Lane de Barros Dantas & Dodd 2021).

To date, studies have explored the relationship between the time of day and genome-wide temperature stress responses in a small number of plant species (Fowler, Cook & Thomashow 2005; Dodd *et al*. 2006; Bieniawska *et al*. 2008; Zhu, Oh, Wang & Wang 2016; Grinevich *et al*. 2019; Li, Gao, Liu, Sun & Tang 2019; Blair *et al*. 2019; Kidokoro *et al*. 2021; Bonnot & Nagel 2021; Bonnot, Blair, Cordingley & Nagel 2021). In Arabidopsis, gating of heat stress responses occurs at both the transcriptome and translatome levels (Bonnot & Nagel 2021). In rice panicles, rhythmic transcripts were observed to be more sensitive to warm nighttime temperatures than those that were nonrhythmic (Desai *et al*. 2021). Furthermore, more recent work showed that in bread wheat the transcriptome response to cold stress is gated, with variations across the three wheat sub-genomes (Graham, Paajanen, Edwards & Dodd 2022). Together suggesting that time of day or gating of temperature stress responses play key roles in the physiological outputs of important crop species. To date the above-mentioned studies have been performed primarily in C3 plants. However, for C4 plants specifically for varieties that are considered more stress tolerant, it is not known whether molecular responses to stress are modulated by the time of day and furthermore whether the occurrence or magnitude of the response varies depending on the sensitivity of specific genetic background.

Previous studies have shown that sorghum (*Sorghum bicolor*), a C4 cereal crop, can tolerate relatively high temperatures compared to other cereals (Sunoj *et al*. 2017; Chiluwal *et al*. 2020). However, sorghum genotypes with different sensitivities to temperature extremes including heat and cold stress have been identified (Chiluwal *et al*. 2020; Ostmeyer *et al*. 2020; Vennapusa *et al*. 2021). Despite, sorghum known to be relatively tolerant to different abiotic stresses, temperature increases above optimum (32°C) can decrease sorghum yields (Prasad, Bheemanahalli & Jagadish 2017; Tack, Lingenfelser & Jagadish 2017). Comparatively, sorghum being a tropical crop is highly sensitive to cold stress particularly during the early season, wherein temperatures below 15°C are known to reduce seedling emergence leading to poor plant stand and lower yields (Kapanigowda *et al*. 2013; Chiluwal *et al*. 2018; Moghimi *et al*. 2019). In summary, sorghum, though known to thrive under adverse conditions, temperatures below or above optimum induces cold and heat stress, respectively, negatively impact the overall physiology and growth.

In sorghum, a large proportion (52%) of the transcriptome shows rhythmic diurnal expression, suggesting critical control by the time of day and/or the clock on cellular processes (Lai *et al*. 2020). A broader understanding of the molecular changes and gene networks that are involved in temperature stress responses is warranted in diverse plant species, including those that are naturally stress-tolerant as this may contribute to a positive outcome in terms of both resilience and yield. In this study, we asked whether there is molecular variation in response to heat and cold stress in thermo-tolerant and thermo-sensitive sorghum genotypes and whether these dynamics are driven by the time of day. For this, we monitored the transcript abundance in heat tolerant (Macia), cold tolerant (SC224), and heat/cold susceptible (RTx430) sorghum varieties at four times of the day (early morning, middle of the day, late afternoon, and 3 h after the beginning of the night) following a 1 h exposure to heat (42°C) and cold (10°C) stress. Our analysis shows a profound control of time of day on the molecular responses to heat and cold stress in sorghum, which can be explained by gene expression rhythmicity. Furthermore, the temperature responses of the thermo-tolerant genotypes are more influenced by time of day, and genes that exhibit significant differences in their response to temperature between the selected genotypes were identified. Most of the genotype effect is observed on the average gene expression, which could explain the different temperature sensitivities. Of significance, non-neglectable numbers of genes exhibited differential temperature responsiveness between thermo-tolerant and sensitive genotypes, including several genes from the Calcium-binding EF-hand family and transcription factors (TFs) that may be ideal targets for genetic manipulation in select varieties.

## Results

### The transcriptome response to temperature stress differs depending on the time of day and the genotype

We first hypothesized that tolerant sorghum may show temporal variation at the molecular level in response to temperature stress. To investigate genotype-specific variation resulting from the time of day and temperature stress, we selected sorghum varieties that have previously been shown to exhibit different temperature sensitivities (Chiluwal *et al*. 2020; Vennapusa *et al*. 2021). Heat and cold-susceptible RTx430, heat-tolerant Macia, and cold-tolerant SC224 were subjected to 1 h heat stress (42°C) or cold stress (10°C) at four different times of the day (ZT1, ZT6, ZT9, and ZT15, Figure 1A), and mRNA-Seq was performed from leaf samples (Supplementary Dataset S1). First, pairwise comparisons revealed that heat stress results in greater perturbation than cold stress regardless of the time of day or genotype (Figure 1B, Supplementary Datasets S2-S3). In total, 2575 and 11218 differentially expressed (FDR < 0.05, |Log_2_ Fold Change| > 1) genes (DEGs) were identified in response to cold and heat stress, respectively. Interestingly, the numbers of differentially expressed genes (DEGs) are very different depending on the time of day, in both experiments (*e.g.* 455 and 130 DEGs up-regulated in RTx430 at ZT1 and ZT15, respectively, Supplementary Dataset S3). In response to heat stress, a large proportion of DEGs are responsive in the morning (ZT1) and middle of the night (ZT15) compared to the afternoon (ZT6) and evening (ZT9, Figure 1B). These observations reflect a greater necessity for the plant to turn on heat-responsive genes when the stress is occurring outside of the time range of naturally occurring high temperatures. Of note, a high number of DEGs is observed at ZT15 under heat stress, especially in RTx430, and we hypothesize that this may be partly explained by a higher temperature change when applying the stress at this time of day, because of a lower control temperature at night (30/20°C day/night). For cold stress, the time of day effect is even more pronounced, with a large subset of DEGs in the morning (ZT1), and a reduced number of DEGs throughout the day (ZT6, ZT9) or the middle of the night (ZT15). Hence, contrary to what was observed in response to heat stress, gene expression was more disturbed when cold stress occurred when temperatures are at their lowest point in natural conditions.

**Figure 1.**
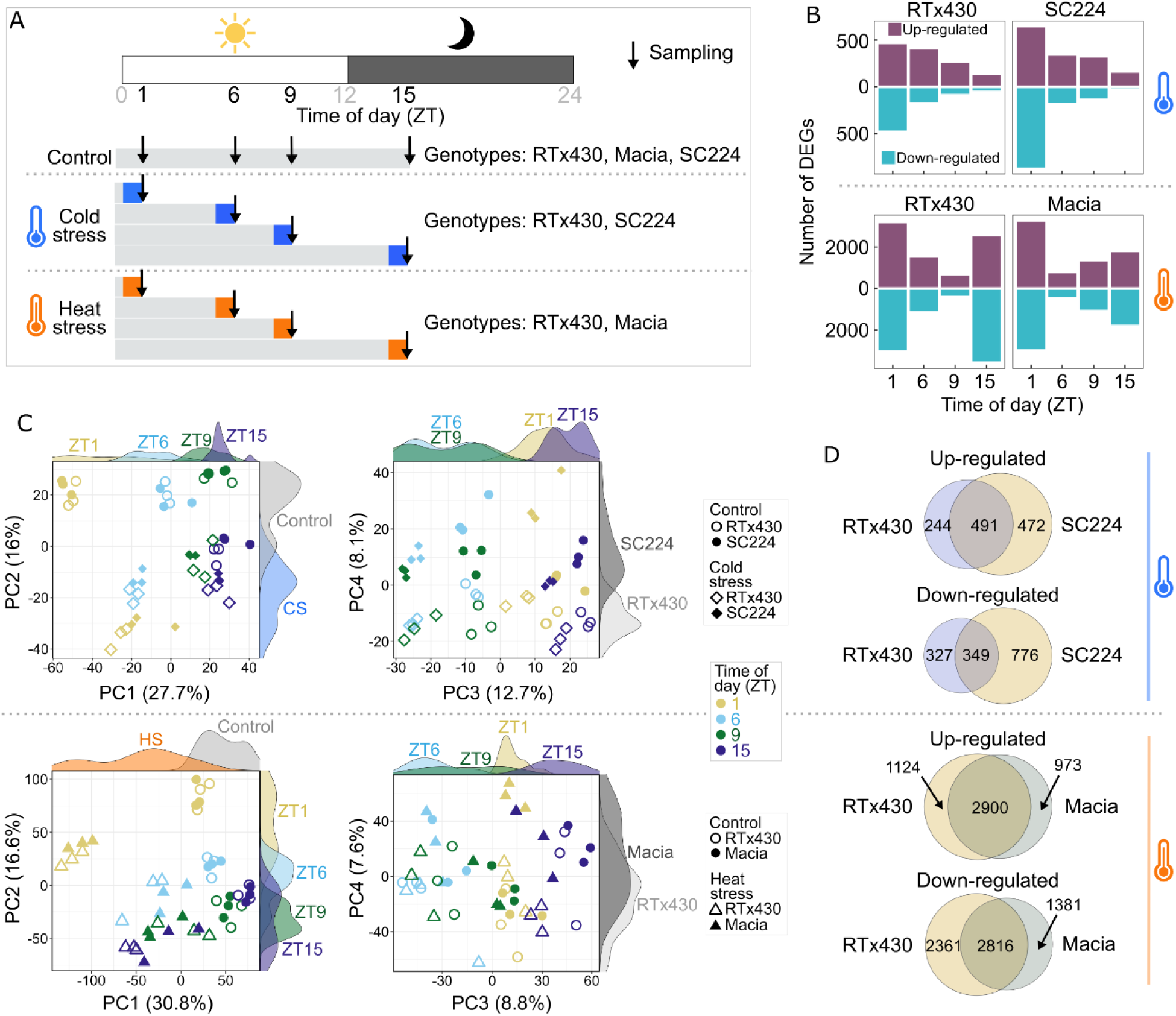
Time of day and genotype influence on the sorghum transcriptome responses to temperature stress. A, Schematic of the experimental design. B, Bar plots representing numbers of DEGs (stress vs control) in response to cold (upper plots) and heat (lower plots) stresses, at different times of day and in different sorghum genotypes. C, Principal component analysis (PCA) of the 2575 and 11218 DEGs identified in the cold stress and heat stress experiments, respectively. Colored areas above and on the right part of each individual PCA plot represent distributions of the indicated groups. CS and HS refer to cold stress and heat stress, respectively. D, Venn diagrams depicting the overlapping DEGs between genotypes, for up-regulated and down-regulated DEGs, and for the cold and heat stress experiments.

Multidimensional analysis performed from expression data of the identified DEGs confirmed the strong influence of temperature and time of day (Figure 1C). Interestingly, PC1 explained 27.7% of the variation of the cold stress data and separated times of day, while time points were separated on PC2 (which explains 16.6% of the variance) for heat stress data (Figure 1C). This suggests a greater influence of time of day on the cold stress-responsive transcriptome than for the heat stress-responsive DEGs. For both experiments, PC4 separates genotypes and explains about ∼8% of the variation (Figure 1C), so we next investigated this effect through a qualitative analysis, by comparing the lists of DEGs between genotypes (Figure 1D). Overall, more genes were differentially expressed in the tolerant (SC224) compared to the sensitive (RTx430) genotypes in response to cold stress. In addition, despite significant overlaps, 49% and 69% of the up-regulated and down-regulated DEGs in SC224 under cold stress were specific to this genotype (Figure 1D). Differences between genotypes for the heat stress experiment seem to be less pronounced, with a higher overlap between DEG lists, especially for up-regulated genes (Figure 1D). However, Macia, which is heat stress tolerant, showed fewer DEGs at ZT6 as compared to RTx430, while fewer genes were impacted at ZT9 in the sensitive RTx430 (Figure 1B). This suggests that time of day contributes to genotype-specific variations in the response to heat stress. Lastly, less DEGs were identified in the heat-tolerant vs susceptible genotype, contrasting with cold stress results (Figure 1D).

All together, these results showed for the first time that the sorghum transcriptome is responding to temperature stress in a remarkable time-of-day specific manner. The influence of time of day is much more pronounced as compared to what we have observed in Arabidopsis at the transcriptome and translatome levels under heat stress (Bonnot & Nagel 2021) when considering the number of DEGs at each individual time point. Nonetheless, this previous study was performed in circadian conditions (*i.e.* absence of environmental cues), whereas photocycles and thermocycles were used in the present study. Consistent with previous studies, we found that C-repeat Binding Factors (CBFs) genes that are known to be transcriptionally activated in response to low temperatures were also significantly induced in response to cold stress in sorghum (Thomashow 1999, Supplementary Figure S1). More generally, TFs are especially well represented within up-regulated DEGs under cold stress (12.4% and 16% in RTx430 and SC224, respectively), as compared to down-regulated DEGs and heat stress-responsive DEGs (5.3% to 7.2%, Supplementary Figure S2). In response to heat stress, we observed significant induction of the Heat Shock Factors (HSFs, Supplementary Figure S3). Interestingly, temperature stress responses of these genes are also gated (*i.e.* different responses depending on the time of day).

### The timing of the response to temperature stress relies on the diurnal gene expression pattern

Both time of day and the circadian clock highly influence gene expression and regulations of abiotic stress responses (Grundy *et al*. 2015; Bonnot *et al*. 2021). Thus, the magnitude of response of a particular gene at a given time point often relies on the rhythm of transcript abundance over the course of the day. To investigate the influence of rhythmic gene expression on our transcriptomic results, we integrated a diurnal transcriptome dataset recently published (Lai *et al*. 2020). This study identified 16,752 (52% of expressed genes) rhythmic gene expression patterns in sorghum, in conditions similar to those used in our present study (Supplementary Dataset S4). A majority of these rhythmic genes showed peak expression in the evening/beginning of the night (Figure 2A). Using this list of rhythmic genes, we first observed that 61 to 71% of our identified DEGs are diurnally regulated under control conditions (Supplementary Figure S4). These numbers, greater than the proportion observed at the genome scale (52%), suggest that thermo-responsive genes tend to be diurnally controlled.

**Figure 2.**
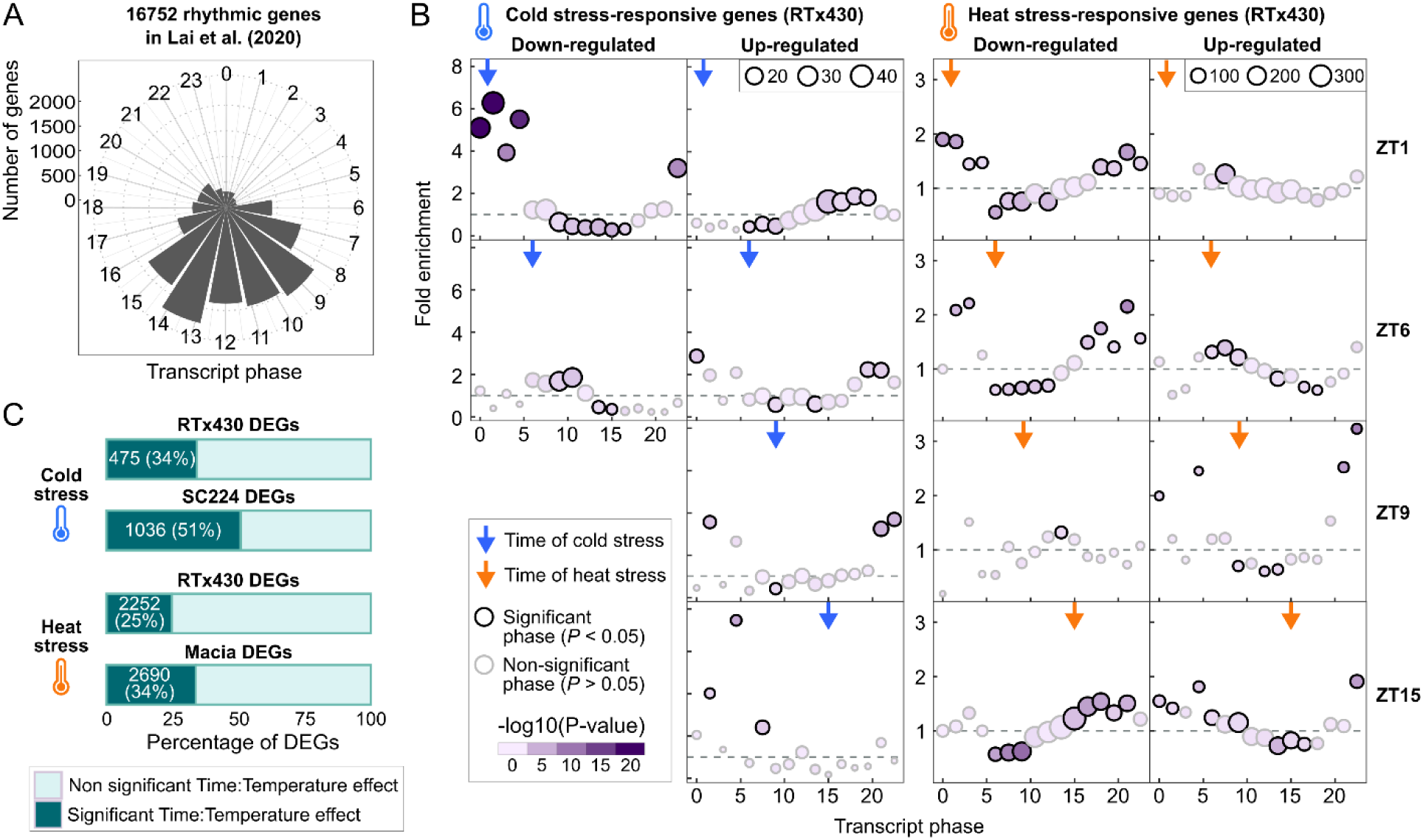
Influence of the rhythmic gene expression pattern on the time of day response to temperature stress. The timing of the response to temperature stress may be related to the diurnal gene expression pattern. A, Circular bar plot representing the counts of the different phases identified in the rhythmic transcriptome from Lai et al., (2020, see the Materials and methods section for details). The phase is defined as the timing of peak abundance (a phase of 0 and 12 indicates a peak abundance at subjective dawn and the beginning of the subjective night, respectively). B, Enriched phases in lists of DEGs presented in Figure 1B. Proportions of the different phases in the lists of DEGs were compared to those of all rhythmic genes identified in Lai et al. (2020). Only genes identified as rhythmic in Lai et al. (2020) wer considered for this analysis. Horizontal gray dashed lines correspond to a fold enrichment of 1. Bubble plots represent over-(fold enrichment > 1) and under-represented (fold enrichment < 1) phases in the list of DEGs as compared to the reference. Chi-Square tests were performed and significance was judged at P-value < 0.05. For a meaningful enrichment calculation, only sets of ≥ 100 DEGs were considered for this analysis. C, Proportions of genes with a significant interaction between the effects of time of day and temperature in lists of DEGs identified in Figure 1C. This analysis was performed for each genotype, and for the cold and heat experiments.

We then hypothesized that thermo-responsive genes peak at particular times during the day. To verify this hypothesis, we compared the proportions of phases (*i.e.* timing of peak expression) in our lists of DEGs (*e.g.* 467 down-regulated DEGs in RTx430 in response to cold stress at ZT1) to the proportions of phases in all 16,752 rhythmic genes identified in Lai *et al* (2020) that were used as the reference. This analysis revealed that when cold stress occurred at ZT1, genes peaking in the early morning are highly over-represented in the list of down-regulated DEGs (Figure 2B, Supplementary Dataset S4). On the contrary, genes with a phase between 15-19.5 (*i.e.* peak of expression at night) are over-represented within up-regulated genes (Figure 2B, Supplementary Dataset S4). More generally, we observed that genes are preferentially down-regulated when temperature stress occurs around their peak of expression, while genes are up-regulated when the stress occurs outside of their timing of peak expression (before or after). This is verified at all time points and for both genotypes in response to cold stress (Figure 2B, Supplementary Figure S5). One interpretation is that genes are expressed at the right time during the day to induce proper responses to changes in environmental stimuli. However, if the stimulus is dramatically changing at an unexpected time of day, the expression of genes acting in the stimulus response need to be adjusted to induce cellular responses. This observation is a little more contrasted for the heat stress experiment, especially at ZT6 and ZT9 (Figure 2B, Supplementary Figure S5). Interestingly, when heat stress hit in the middle of the light period (ZT6), genes peaking at that time were over-represented within up-regulated genes. Several genes correspond to HSFs and Heat Shock Proteins (HSPs, Supplementary Dataset S4). Despite an obvious influence of the rhythmic gene expression pattern on the timing of the response to temperature stress, potential phase differences exist between our selected genotypes, which cannot be resolved with our datasets. In addition, the diurnal rhythmic transcriptome has been identified in a different genotype BTx623 (Lai *et al*. 2020).

Although expression rhythmicity can explain different magnitudes of a response to temperature stress, genes with constant expression throughout the day could respond in a time-of-day specific manner. To identify all sorghum genes with responses to cold and heat stress that depend on time of day (either specific or with different magnitudes of response), we performed a statistical analysis considering all time points and looking at the interaction between the effects of temperature and time of day (Supplementary Dataset S5). This analysis revealed that cold-responsive DEGs are, in proportion, more influenced by time of day than heat-responsive DEGs (34-51% vs 25-34%, Figure 2C). This supports the observations from the multi-dimensional analysis above (Figure 1C). Furthermore, the response of 51% of the cold stress-responsive DEGs in SC224 was affected by time of day (Figure 2C), while the proportion is 34% in RTx430. Similarly, 34% and 25% of the DEGs in Macia and RTx430 respond to heat stress in a time of day specific manner, respectively (Figure 2C). Of significance, the temperature responses of the thermo-tolerant genotypes are more influenced by time of day. We anticipate that this observation could reflect a potential greater control of the temperature responsiveness by the circadian clock in these lines. Clock genes, however, did not show obvious differences between genotypes in their temperature stress responsiveness (Supplementary Figure S6, Supplementary Table S1).

### Differences between genotypes are mostly explained by different average expression levels

Comparing the lists of DEGs between genotypes allowed for a simplistic visualization of specificity in the temperature responsiveness of thermo-tolerant vs sensitive genotypes (Figure 1D). To get a better estimation of the number of genes with genotypic variations in gene expression, we performed a statistical analysis considering the whole dataset (two separate analyses for the cold and heat stress experiments) and looked for genotype effects, in interaction or not with the temperature and time of day effects (Supplementary Dataset S5, see methods for details). In total, the expression of 5024 and 5989 genes were significantly (FDR < 0.05) affected by genotype effect, in the cold and heat stress experiments, respectively (Figure 3A). From data obtained in control conditions for the 5024 genes, hierarchical clustering revealed four groups with distinct profiles (Figure 3A). Interestingly, these groups showed different levels of gene expression between RTx430 and SC224 (Figure 3A). Including the data obtained under cold stress does not alter this observation, suggesting that this is independent of temperature (Figure 3A, violin plots). Comparable results were obtained in the heat stress experiments (Figure 3A). However, in the six identified transcript groups, temperature differences can be detected (Figure 3A, violin plots).

**Figure 3.**
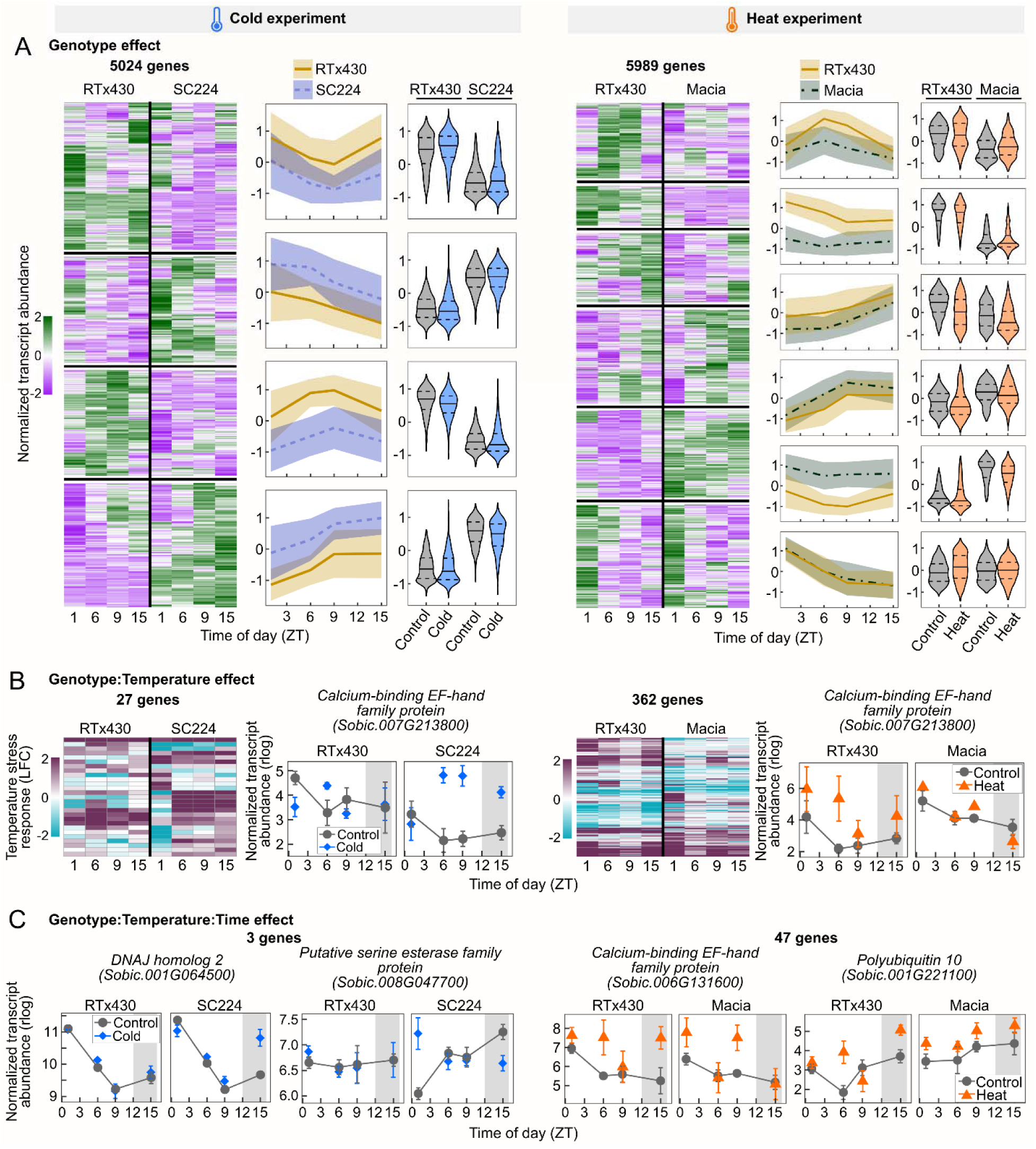
Genotype influence on gene expression levels and temperature stress responses. A, Representation of genes with a significant effect of the genotype (5024 and 5989 DEGs in the cold and heat experiments, respectively). Heatmaps represent the normalized transcript abundance in the control conditions for the genes with a genotype effect. Data are scaled by row and are means of *n* = 3 biological replicates. The transcript abundance over time into four and six groups identified from these heatmaps are represented next to each heatmap, in the cold and heat stress experiments, respectively. On these line plots, solid and dashed lines represent the mean and standard deviation, respectively. Violin plots represent the distributions of normalized transcript abundances within each group, in control conditions and in response to stress, for the two studied genotypes. On these violin plots, solid lines represent medians and the two dashed lines represent the 1st and 3rd quartiles. B, Representation of genes with a significant interaction between the effects of genotype and temperature (27 and 362 DEGs in the cold and heat experiments, respectively). Heatmaps represent the LFC (stress vs control) values, blue and purple indicating a down-regulation and an up-regulation, respectively. For each experiment, transcript abundance profiles of a selected gene are shown (data ± SD, *n* = 3). C, Transcript abundance profiles of selected genes with a significant interaction between the effects of genotype, temperature, and time of day (3 and 47 DEGs in the cold and heat experiments, respectively).

Thus, we next investigated the genotype:temperature effect, and found 27 and 362 significant (FDR < 0.05) genes in response to cold and heat stress, respectively (Figure 3B, Supplementary Datasets S5-S6). The difference in the number of significant genes between experiments is not surprising given the much higher number of DEGs identified in response to heat stress as compared to cold stress (Figure 1). Interestingly, several (five and four in the cold and heat datasets, respectively) genes found by this analysis code for calcium-binding EF-hand family proteins (Supplementary Datasets S5-S6). For example, *Sobic.007G213800* was revealed in both datasets and showed significant differences in its response to temperature stress between the selected genotypes (Figure 3B). This gene showed *i*) different gene expression levels between genotypes under control temperature (RTx430 > SC224 and Macia > RTx430) and *ii*) greater response to stress in specific genotypes (SC224 > RTx430 and RTx430 > Macia).

To further study the influence of time of day on genotype-specific temperature responses, we selected genes with a significant genotype:temperature:time effect and identified three and 47 genes in the cold and heat stress experiments, respectively (Figure 3C, Supplementary Datasets S5-S6). The selected genes, therefore, respond to temperature stress in a time of day and genotype-specific manner. These highly specific responsive genes are involved in diverse processes related to signaling, and metabolism and include transcriptional regulators (Figure 3C, Supplementary Datasets S5-S6).

Altogether, these results demonstrate that few genes exhibit significant differences in their response to temperature between the selected genotypes. Most of the genotype effect is observed on the average gene expression, which could explain the different temperature sensitivities. Nonetheless, a subset of genes exhibited differential temperature responsiveness between thermo-tolerant and sensitive genotypes, including several genes from the calcium-binding EF-hand family and TFs. These regulatory genes might contribute to genotype-specific fine-tuning of thermo-responsive pathways. In addition, the presence of kinases and members of the RING/U-box family suggests specificities at other regulatory levels such as post-translational regulation.

### Gene co-expression network analysis identifies modules regulated by temperature stress in a time-of-day and genotype-specific manner

To identify genes with specific patterns of expression and response to temperature, we employed a weighted gene co-expression network approach. For this analysis, we considered the whole dataset and not only our identified DEGs, after the removal of lowly expressed genes (see methods and Supplementary Figure S7 for details). Four co-expression networks were built, one per genotype and experiment (Supplementary Figures S8-S11, Supplementary Dataset S7). We first observed that more modules (*i.e.* groups of gene nodes with similar expression patterns) were identified in the thermo-tolerant as compared to the sensitive genotypes (29 vs 18 and 20 vs 17 for the cold and heat stress experiments, respectively, Figure 4). This larger diversity in transcript accumulation profiles in thermo-tolerant genotypes could reflect a finer transcriptional regulation under temperature stress. However, similarities between modules identified in the two genotypes make the identification of highly genotype-specific patterns difficult (Supplementary Figure S8-S11). This is not unexpected given the relatively low numbers of genes with a significant interaction between the effect of temperature and genotype (Figure 3B).

**Figure 4.**
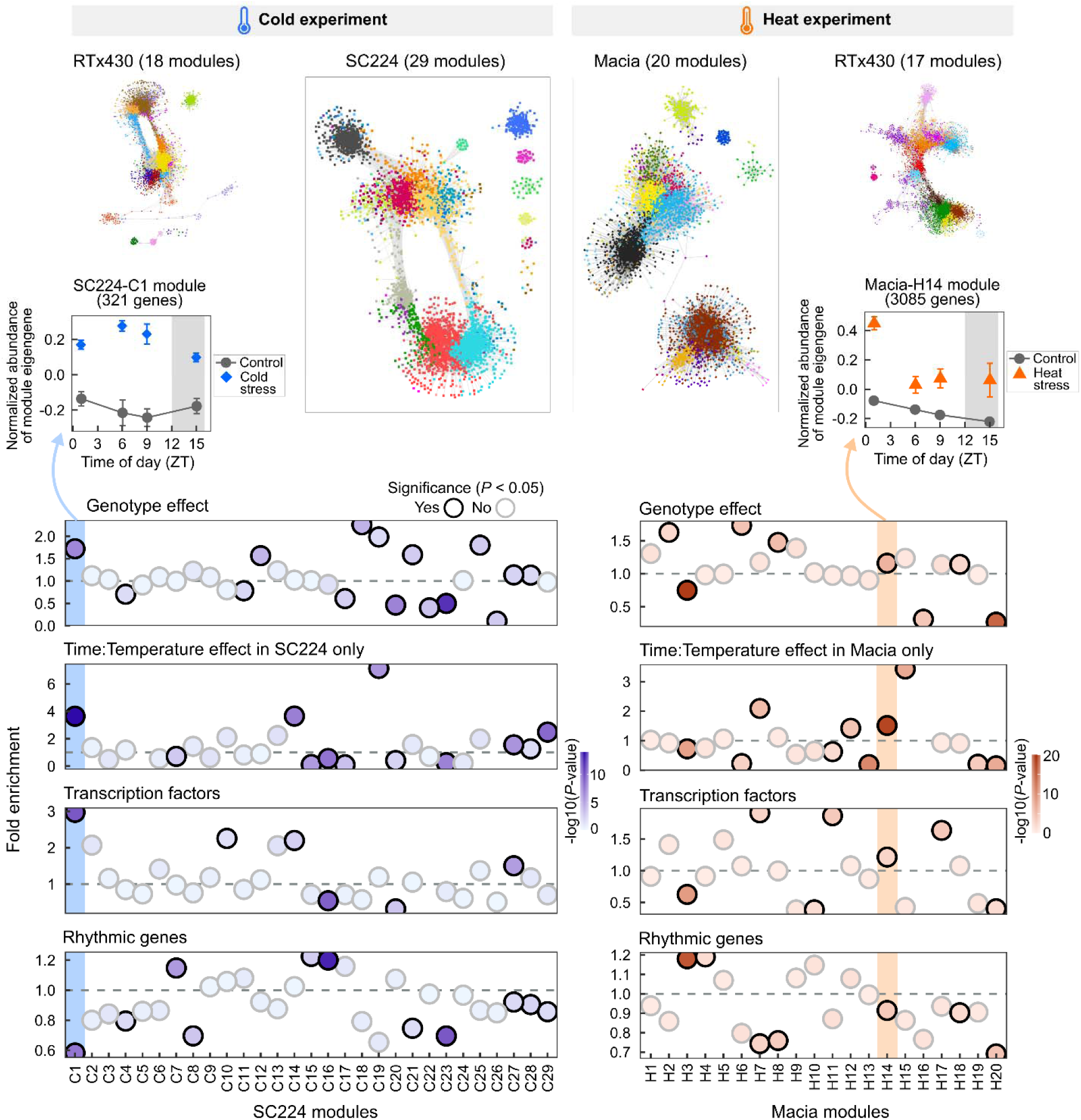
Identification of gene modules with temporal and genotype-specific responses to temperature stress using a co-expression network approach. Network visualization was done in Cytoscape using a Prefuse Force Directed layout, with an edge threshold cutoff of weight > 0.15. Gene nodes are colored by module membership. Colors do not reflect similarities between networks and were randomly attributed for each network analysis. Bubble plots represent the enrichment of specific lists of genes within modules identified in SC224 and Macia in the cold stress and heat stress experiments, respectively. Genes with a genotype effect and a time:temperature effect specific to either SC224 or Macia were represented in Figures 3A and 2C, respectively. TFs were identified from PlantTFDB. Rhythmic genes were described in Lai et al. (2020). Fold enrichment < 1 and > 1 correspond to an under-and over-representation in the module, respectively. Profiles of the module eigengene for modules SC224-C1 and Macia-H14 are highlighted. Gray areas represent the night period. Profiles of all module eigengenes are represented in Supplementary Figure S8-S11.

Thus, to identify gene modules representing genotype specificities, we looked for genes with *i*) a significant genotype effect (highlighted in Figure 3A) and *ii*) a temperature stress response controlled by time of day (highlighted in Figure 2C) in the thermo-tolerant genotypes only, within each individual gene module revealed in the tolerant genotypes (Figure 4). In addition, we searched for TFs and rhythmic genes. This analysis allowed us to reveal modules SC224-C1 and Macia-H14 in the cold and heat stress datasets, respectively (Figure 4). SC224-C1 groups 321 genes that are highly up-regulated under cold stress, with a greater induction (on average) at ZT6 and ZT9. Within this module, genes with a significant genotype effect and/or a cold stress response gated by time of day specifically in SC224 are over-represented (Figure 4). Biological processes related to transcriptional regulation are significantly enriched, confirmed by an over-representation of TFs within this module (Figure 4). The ERF TF family is the most represented, including three members of the CBF subfamily (Supplementary Figure S12). Similar characteristics were found for Macia-H14 from the heat stress dataset (Figure 4). This large module of 3085 genes is highly induced under heat stress, with a greater induction in the early morning (ZT1). Also enriched for TFs, this module contains 13 members of the HSF family (Figure 4, Supplementary Figure S12). In addition, the enrichment of GO terms ‘unfolded protein binding’, ‘chaperone binding’ and ‘nucleus’ suggests a role for this module in the activation of HSF-dependent thermal responses (Supplementary Table S2). The two modules SC224-C1 and Macia-H14, therefore, represent interesting regulatory modules impacted by temperature stress in a time of day and genotype-specific manner.

Surprisingly, we observed a significant under-representation of rhythmic genes within these modules (Figure 4). We previously discussed the strong influence of gene expression rhythmicity on their diurnal gating responsiveness to temperature stress (Figure 2). In addition, the high proportion of rhythmic genes within our lists of DEGs suggested to us that most stress-responsive genes would be diurnally controlled. This new result does not question these conclusions but emphasizes that a significant proportion of genes with a strong thermal response are controlled by time of day under temperature stress conditions only. For example, this is the case for *Sobic.002G269300* (*CBF3*) under cold stress, *Sobic.003G039400* (*HSP17.6*) under heat stress, as well as several members of the Ca^2+^-binding EF-hand family.

### Time of day influences gene rhythmicity and temperature stress responses of EF-hand gene family members

During our analyses of the influence of genotype on the transcriptome response to temperature described above, several genes encoding EF-hand Ca^2+^-binding proteins were revealed (Figure 3). Calcium-binding EF-hand-containing genes belong to a family of proteins that function as Ca^2+^ sensors (Day, Reddy, Shad Ali & Reddy 2002; Mohanta, Kumar & Bae 2017). As Ca^2+^ is an important cellular messenger that plays a role in responses to hormones and external stresses, for example, we speculate that these proteins may contribute to genotype-specific thermo-tolerance in sorghum (Reddy, Ali, Celesnik & Day 2011). We identified 161 members within this family that contain known protein domains (see Methods, Supplementary Dataset S8). The expression of about half was influenced by genotype, either in the cold stress or heat stress experiment, or both (Figure 5A). A more detailed analysis showed that 37.3% of the family had a significant genotype effect in the heat stress experiment and that this proportion was significantly higher than that of all analyzed genes (23.5%, Figure 5B). Despite a similar trend in the cold-stress experiment, no significant enrichment was observed, suggesting a more specific difference in gene expression between Macia and RTx430 (heat-stress experiment) than between SC224 and RTx430 (cold-stress experiment), for this gene family. However, of the 27 and 362 genes with a significant Genotype:Temperature effect in the cold and heat stress experiments, respectively (Figure 3B), six (22%) and nine (2.5%) corresponded to EF-hand Ca^2+^-binding proteins and therefore showed differential temperature responsiveness between thermo-tolerant and sensitive genotypes.

**Figure 5.**
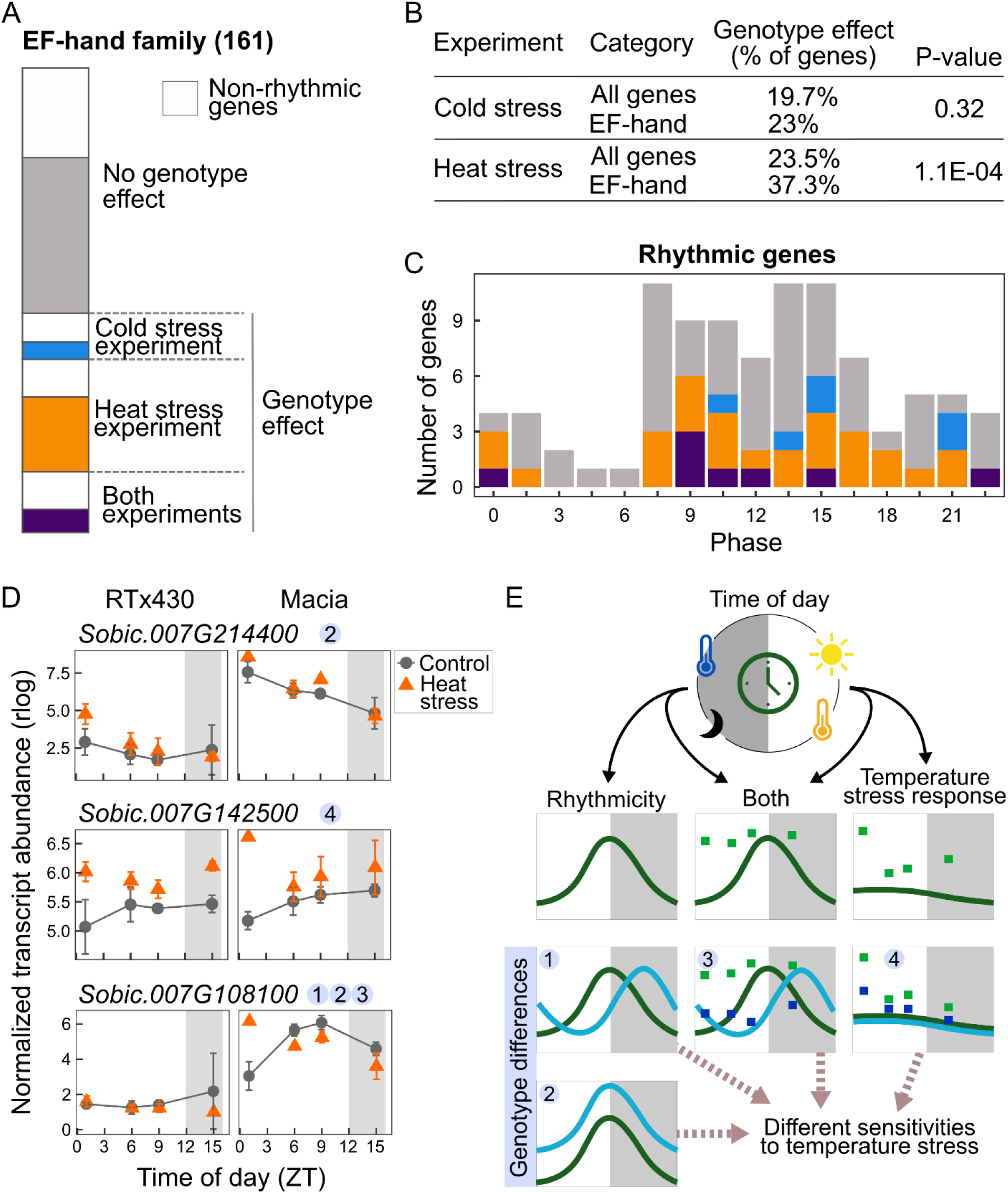
Time of day influences gene rhythmicity and temperature stress responses: Example shown for EF-hand gene family members. A, Stacked barplot representing the proportion of genes having or not a genotype effect in the cold stress and heat stress experiments (shown in Figure 3A), from 161 members of the EF-hand family. B, Table indicating the proportions of genes with a genotype effect within all expressed genes and EF-hand family members highlighted in A. Fisher’s exact tests were performed to compare the proportions between the two groups. C, Phase (i.e. timing of peak expression) distribution of the rhythmic EF-hand family members highlighted in A. Rhythmic genes and phases were identified in Lai et al. (2020, see Materials and Methods for details). D, Transcript abundance profiles of selected EF-hand family members (means ± SD, *n* = 3). E, Schematic representation of the influence of time of day and genotype on the gene expression pattern and response to temperature stresses. In D and E, grey areas represent the night period.

We next looked at the timing of gene expression of the EF-hand Ca^2+^-binding gene family and observed that most members with a rhythmic expression peak in the evening and early night (ZT7.5 – ZT15, Figure 5C), as observed for all 16752 rhythmic genes identified in sorghum (Figure 2A, Lai et al., 2020). No specific phase distribution was identified for subsets of genes affected or not by the genotype effect (Figure 5C). This suggests that the EF-hand Ca^2+^-binding family is acting at multiple times of day, and that genotypic variability in gene expression is not more pronounced for members with specific expression patterns. Regardless of whether members of this family exhibit a rhythmic profile, their expression is influenced by genotypes in different ways, as illustrated with selected genes in Figure 5D. For example, although *Sobic.007G214400* did not show high magnitude of response to heat stress, its expression level is significantly higher in heat-tolerant genotype Macia as compared to RTx430 (Figure 5D). On the contrary, *Sobic.007G142500* showed similar expression levels between the two genotypes but exhibited a stronger response to heat stress in Macia when the stress occurred in the early morning (ZT1, Figure 5D). Interestingly, *Sobic.007G108100* showed very different expression patterns between genotypes, with a higher and different pattern of expression in Macia, and a specific up-regulation under heat stress in the morning (ZT1), in Macia only (Figure 5D).

In this study, our results revealed that both temperature stress and gene expression rhythmicity highly affect changes in transcript levels in sorghum (Figure 5E).

As illustrated with the EF-hand Ca^2+^-binding protein family, genotype differences are observed either on the average gene expression level, or on the temperature stress response, either in interaction with time of day or not. We speculate that these genotype differences, summarized in Figure 5E, may lead to different sensitivities to temperature stress.

## Discussion

Temperature stress can limit yield potential in many crops including sorghum (Zhang, Zhao & Zhu 2020). Interaction with the environment is tightly coupled to molecular dynamics and cellular stress responses in plants. Here we report for the first time that the time of day modulates or gates the molecular response to temperature stress in sorghum, a C4 crop, and this is further refined by the genotype sensitivity (Figures 1, 2, and 5). In general, genes with gated responses to temperature stress show either *i*) a specific response to stress at a particular time of day (and no significant change in gene expression at other time points), *ii*) an opposite response to stress between time points (*e.g*. up-regulated at a given time of day, and down-regulated at another time point), or *iii*) a similar response (up-regulated or down-regulated) at multiple times of day, but with different magnitudes of response depending on the time of day.

Major aspects of the time of day regulation of transcript abundance in response to temperature stress are controlled by the circadian clock or exhibit some form of diurnal expression (Bonnot *et al*. 2021). Our analysis revealed that up to 70% of our identified DEGs are diurnally regulated (Supplementary Figure S4). These numbers are greater than the proportion observed at the genome scale (52%) supporting that thermo-responsive genes tend to be diurnally controlled (Covington *et al*. 2008). However, it is worth noting that the gating of oscillator components to either heat or cold stress is not as evident as other rhythmic genes (Supplementary Figure S6). Of the sorghum clock genes, *Sb_PRR73* (*Sorghum bicolor_Pseudo Response Regulator 73*) shows a gated response to both heat (up-regulated) and cold stress (down-regulated), with a specific response in the morning (ZT0). *Sb_GI (Sorghum bicolor_Gigantea)* is downregulated in response to cold only at ZT0 and upregulated in response to heat at ZT0 and ZT15 and interestingly the response to both heat and cold is more pronounced in RTx430, the sensitive genotype (Supplementary Figure S6). In terms of heat stress, the response for *Sb_GI* is similar to what was observed in Arabidopsis where the increased transcript accumulation occurs before (morning) and after (subjective dark) the peak of *GI*’s expression (Bonnot & Nagel 2021). In a recent study in wheat, *TaPRR73* and *TaGI* are also the two oscillator components that show strong perturbation in response to cold stress resulting in a delay in their peak of expression (Graham *et al*. 2022). These observations suggest that the sensitivity to temperature for some oscillator components may be conserved across species while others may not be. GI and members of the PRRs in sorghum play key roles in flowering and thus circadian gating of their molecular response to heat stress, for example, may be directly related to flowering time changes (Murphy *et al*. 2011; AbdulLAwal, Chen, Xin & Harmon 2020). Alternatively, different circadian signaling components may separately regulate expression rhythmicity under control conditions and gate stress responsiveness depending on the plant as suggested by Graham et al. (2022) for cold responses in wheat.

In Arabidopsis, we previously showed that time of day gates the heat stress response of about a third of the circadian-regulated heat stress-responsive transcriptome and translatome (Bonnot & Nagel 2021). In sorghum, similar proportions (25-34%) of the heat-responsive transcriptome are gated by the time of day, depending on the genotype (Figure 2C). The genome-wide gating response to temperature is not restricted to heat stress and was also observed under cold stress, as previously reported in Arabidopsis (Blair *et al*. 2019), and recently in wheat (Graham *et al*. 2022). Our observations along with published work raise the intriguing question of whether specific clock components gate the molecular dynamics in response to heat stress or cold stress or both. Temperature gating experiments in multiple circadian clock mutants may help to shed light on this. Furthermore, time of day control on temperature stress response was primarily performed in whole seedlings or plants. Work in Arabidopsis suggests that different parts of the plant show variation in circadian rhythms and that multiple points of clock coordination may exist (Gould *et al*. 2018). However, it is not known the extent of time of day control in specific tissues and/or cell types in response to stress but this might help to further dissect the regulatory mechanism of gating and its contribution to physiological responses. It is also worth noting that previous work has shown that the circadian clock runs slower in cultivated tomatoes than in their wild relatives suggesting one aspect of internal clock adjustment that may have adapted to specific geographic location and environment (Müller *et al*. 2016). It would be worthwhile to examine whether circadian gating in response to stress is conserved or adjusted in these lines and similar varieties.

Interestingly, our data indicate that the influence of time of day on the transcriptome response is more pronounced in thermo-tolerant varieties, which might reflect more robust clock control of molecular changes or less sensitivity in stress perception. Future global analysis of gene expression rhythmicity in diurnal and circadian conditions, in multiple sorghum genotypes with different sensitivities to temperature stress, is necessary to unravel the precise mechanism. Nonetheless, the gating response to temperature cannot be fully explained by gene expression rhythmicity. Indeed, our analyses revealed modules of genes whose temperature stress response is strongly gated by time of day, in which rhythmic genes are significantly under-represented (Figure 5). This result evidenced that non-rhythmic genes are also subjected to gating of temperature responses and may serve as an alternative or backup regulatory mechanism for the plant to deal with unpredictable environmental changes.

In our study, we identified potential target genes that can be used to improve crop thermo-tolerance based on their genotype specificities. Our analyses showed that most genes with a genotypic effect showed a greater average expression level in a particular genotype as compared to the other, independently of the temperature condition. Some genes, such as *Sobic.004G108100* – a member of the Ca^2+^-binding EF-hand protein family – also respond to heat stress in a time-of-day and genotype-specific manner (Figure 5). Cycles of Ca^2+^ are observed in the cytoplasm during the day and are controlled by both the circadian clock and light signaling (Xu *et al*. 2007; Martí Ruiz, Jung & Webb 2020). Oscillations of cytosolic free Ca^2+^ also regulate circadian clock function through a mechanism involving the Ca^2+^-sensor CALMODULIN-LIKE24 (Martí Ruiz *et al*. 2018). Under abiotic stresses, intracellular Ca^2+^ concentration increases, Ca^2+^ playing a role in both the sensing of stress and in signal transduction through downstream Ca^2+^-binding proteins (Dong, Wallrad, Almutairi & Kudla 2022; Li, Liu, Jin & Peng 2022; Xu, Niu & Jiang 2022). In maize, under saline-alkaline stress, Ca^2+^ binds to ZmNSA1 – a Ca^2+^-binding EF-hand protein – triggering its degradation, which promotes root Na+ efflux and ultimately, saline-alkaline tolerance (Cao *et al*. 2020). In rice, the overexpression of the annexin *OsANN1* improves plant growth under abiotic stress (Qiao *et al*. 2015). Ca^2+^-binding proteins, therefore, represent interesting targets for genetic improvement.

## Conclusions

Improving the resilience of sorghum varieties to temperature changes relies on a comprehensive understanding of the molecular basis of thermotolerance or susceptibility. Genes with genotype-specific regulation could also represent candidate markers of differential sensitivity to temperature stress as highlighted above. Using field trials with multiple sorghum genotypes, transcript levels could be measured for the identified candidate markers. Transcript levels and genetic markers could further be used in models for predicting important crop traits and thermo-tolerance of specific genotypes (Azodi, Pardo, VanBuren, de los Campos & Shiu 2020).

## Materials and Methods

### Plant Materials and Growth Conditions

The sorghum genotypes used in this study are RTx430 (cold and heat-sensitive inbred line), Macia (heat-tolerant inbred), and SC224 (cold-tolerant inbred) (Chiluwal *et al*. 2020; Vennapusa *et al*. 2021). Plants were grown in controlled environment chambers (Conviron model PGR15; Winnipeg, MB, Canada) under control (30/20°C; maximum day/night temperatures) conditions. The chambers were programmed to reach the daytime (0800 to 1700 h) target temperature of 30°C, by following a gradual increase from 20 to 30°C (control) with a 3 h transition (0500 to 0800 h). Similarly, the nighttime (2000 to 0500 h) target temperature of 20°C was obtained by a gradual decrease in temperature from 30 to 20 °C with a 3h transition (0500 to 0800h). The chambers were maintained at 12 h photoperiod, with 800 μmol m^-2^ sec^-1^ light intensity at 5 cm above the canopy and 60 % relative humidity (RH). After 7 days of seedling emergence the seedlings were subjected to 1 h of cold (10°C) or heat stress (42°C) at four different times of the day (ZT0, ZT6, ZT9, and ZT15; Figure 1A), wherein 0 h is when the lights were switched on inside the chamber (considered as onset of dawn). Seedlings from controlled environment chambers were moved to either cold or heat stress chambers at 0, 5, 8 and 14 h after dawn and exposed to either cold or heat stress for an hour before sampling. The seedlings were sampled after the stress period of 1 h and immediately frozen in liquid nitrogen and stored at −80°C.

### mRNA Isolation, Sequencing, and data processing

Between ∼2 and ∼40 ug of total RNA was extracted from three biological replicates of sorghum samples for each genotype and treatment (control, heat, cold stress) using GeneJET Plant RNA Extraction Kit (Thermofisher K0801) followed by DNAse I treatment (Thermofisher EN0521). Total mRNAs were isolated using biotinylated oligo(dT) and streptavidin magnetic beads (New England Biolabs) as previously described (Wang *et al*. 2011). Purified mRNAs were used for library preparation as previously described with the following modifications. In the final enrichment step, indexed adapter enrichment primers were used (Townsley, Covington, Ichihashi, Zumstein & Sinha 2015) and 12 cycles were performed to amplify the libraries. The libraries were quantified using a Qubit 2.0 Fluorescence Reader (Thermo Fisher Scientific) and quality was verified using a Bioanalyzer 2100 (Agilent Genomics). Final libraries were multiplexed and sequenced on the NextSeq 500 (Illumina) at the UC Riverside (UCR) Institute for Integrated Genome Biology (IIGB) Genomics Core facility to obtain 75 nt single-end reads. Sequencing reads were not trimmed and mapped on the Sbicolor_454_v3.1.1 genome using Hisat2 and the SystemPipeR pipeline (genome downloaded from Phytozome: https://data.jgi.doe.gov/refinedownload/phytozome?organism=Sbicolor&expanded=45). Read counting was performed with the summarizeOverlaps function from the GenomicRanges Package, and using the Sbicolor_454_v3.1.1.gene.gff3.gz file for the annotation. Normalized transcript abundance was then calculated using the rlog function from the R package ‘DESeq2’(Love, Huber & Anders 2014), and is provided in Supplementary Dataset S1.

### Statistical analyses

All statistical analyses were performed with the software program R v 4.1.2 (R Core Team, 2022). Pairwise comparisons were used to compare temperature stress vs control conditions at each time of day and for each genotype, and were performed on raw counts with the R package ‘DESeq2’, and using the SystemPipeR workflow (Love *et al*. 2014; Backman & Girke 2016). Significant differences were based on an FDR < 0.05 and Log_2_ Fold Change > |1|. Results are provided in Supplementary Dataset S2. Phase enrichment analysis was performed using the tool ‘CAST-R’ (Bonnot, Gillard & Nagel 2022). All genes with rhythmic expression identified in Lai et al. (Lai *et al*. 2020) were used as the reference. Phase values correspond to LAG values in the JTK_Cycle output, that were adjusted to circadian time (CT) with the following calculation: CT phase = (JTK_Cycle LAG/estimated period) * 24 (mentioned as CT.PHASE in Lai et al., 2020) To reduce the number of phase groups identified in Lai et al. (2020), phases were rounded as follows: [0, 0.75] = 0; (0.75, 2.25] = 1.5; (2.25, 3.75] = 3; (3.75, 5.25] = 4.5; (5.25, 6.75] = 6; (6.75, 8.25] = 7.5; (8.25, 9.75] = 9; (9.75, 11.25] = 10.5; (11.25, 12.75] = 12; (12.75, 14.25] = 13.5; (14.25, 15.75] = 15; (15.75, 17.25] = 16.5; (17.25, 18.75] = 18, (18.75, 20.25] = 19.5; (20.25, 21.75] = 21; (21.75, 23.25] = 22.5. Briefly, under-represented (phase enrichment < 1) and over-represented (phase enrichment > 1) phases in the selected subset of genes (e.g. down-regulated DEGs at ZT1 under cold stress in RTx430) are identified by comparing the proportions of each individual phase within the subset of genes with those of the reference. Significant differences are assessed at *P* < 0.05 using Chi-squared tests. Only genes identified as rhythmic in Lai et al. (2020) were considered for this analysis. As suggested in the ‘CAST-R’ application, phase enrichment was performed on subsets of genes with a minimal list size of 100 genes. Results of the phase enrichment analyses are provided in Supplementary Dataset S4.

To analyze the interaction between the effects of temperature and time of day, likelihood ratio tests (LRT) were performed using the ‘DESeq2’ package (Love *et al*. 2014). Four LRTs were performed, one per experiment (cold stress and heat stress) and genotype. LRTs are conceptually similar to an analysis of variance (ANOVA) calculation in linear regression (Love et al., 2014). The full model was as follows: design = Expression ∼ Time + Temperature + Time:Temperature. For this analysis, only genes with total read counts > 10 were considered (25,532 remaining genes). To analyze the interaction between the effects of temperature and genotype, and between temperature, genotype, and time of day, two LRTs were performed (one for each temperature stress experiment), using the following model: design = Expression ∼ Time + Temperature + Genotype + Time:Temperature + Time:Genotype + Temperature:Genotype + Time:Temperature:Genotype. Statistical results are provided in Supplementary Dataset S5.

To identify the enrichment of rhythmic genes, TFs, genes with a significant genotype effect, and/or genes with a significant interaction between the effects of temperature and time of day within the network modules, Fisher’s exact tests were performed. Proportions of these specific lists of genes within each individual module were compared to those in all genes present in the network analysis. Significant differences were judged at *P* < 0.05. Gene Ontology (GO) enrichment analysis was performed with agriGO v 2.0 (Tian *et al*. 2017). GO terms with a fold enrichment > 1 and FDR < 0.05 were considered as significantly over-represented in the selected subset of genes.

### Data mining and visualization

Principal Component Analysis (PCA) was performed from normalized (rlog) expression data of identified DEGs, using the multivariate data analysis R package ‘ade4’ (Thioulouse, Chessel, DoléDec & Olivier 1997). Venn diagrams were performed using the R package ‘VennDiagram’ (Chen & Boutros 2011). Heatmaps were generated using the R package ‘pheatmap’ (Kolde, 2019). Within heatmaps, the number of clusters (groups of genes with similar expression patterns) was determined manually. Bar plots, violin plots, line charts, and bubble plots were visualized using the R package ‘ggplot2’ (Wickham, 2016).

Weighted Gene Co-expression Network Analysis was performed using the R package ‘WGCNA’ (Langfelder & Horvath 2008). To remove genes that introduce noise into the network analysis, only genes with read counts > 10 in at least 50% of the samples were considered. The analysis was then performed from normalized expression (rlog) values. Four independent signed networks were constructed, one per experiment (cold stress and heat stress) and genotype. Adjacency matrices were built using a soft threshold power of 18. To identify network modules, a minimum module size of 30 was used, and similar modules were merged using a dissimilarity threshold of 0.25. Networks were visualized with the CYTOSCAPE software v 3.9.0 (Smoot, Ono, Ruscheinski, Wang & Ideker 2011), using a Prefuse Force Directed layout and an edge threshold cutoff of weight > 0.15. Module eigengene values were used to visualize the module expression patterns, and are provided in Supplementary Dataset S7.

To facilitate the visualization of expression patterns of individual genes, an application has been built, with the R package ‘Shiny’ (Chang et al., 2020, see Data Availability section).

### Identification of specific genes and gene families

Circadian clock genes were identified in Lai et al. (2020). Lists of TF families were downloaded from PlantTFDB v 5.0 (Jin *et al*. 2017). Members of the CBF subfamily in sorghum were identified from the annotation of Arabidopsis best hits, provided in the sorghum v 3.1.1 annotation file, downloaded from Phytozome (https://data.jgi.doe.gov/refinedownload/phytozome?organism=Sbicolor&expanded=45). Orthologous genes of sorghum and maize were identified in (Xianjun Lai, Yan & Schnable 2017). Expression data of specific TF families in maize were downloaded from (Li *et al*. 2020). Members of the EF hand Ca^2+^-binding protein family were identified by searching for specific protein domains (PF00036, PTHR10891, PS00018, PS50222, SM00054, and SSF47473) in sorghum protein sequences, using PhytoMine, implemented by Phytozome (https://phytozome-next.jgi.doe.gov/phytomine/). The list of identified sorghum members of the EF-hand Ca^2+^-binding protein family is provided in Supplementary Dataset S8.

## Data Availability

All data reported in this manuscript are accessible from the Gene Expression Omnibus (GEO) database, www.ncbi.nlm.nih.gov/geo (Accession no. GSE225632). The application to visualize results for individual genes can be accessed at https://nagellab.shinyapps.io/Sorghum_time_stress/. R scripts used to process and analyze the data can be accessed at https://github.com/Nagel-lab/Sorghum_temperature_stress.

## Acknowledgments

We would like to thank Amaranath Vennapusa and Nisaraga Narayana for their help during sample collection. We thank Clay Clark and Holly Clark for their expertise in library preparation and sequencing, and Dr. Daniela Cassol for assistance in the use of the SystemPipeR workflow. Funding: This work was partial supported by NSF Early Career Award IOS 1942949 to D.H.N.

## Author Contribution

D.H.N. conceived the project, T.B., I.S., S.V.K.J, and D.H.N. designed and performed experiments. T.B. and D.H.N. analyzed the data and wrote the manuscript. T.B., I.S., S.V.K.J, and D.H.N revised and finalized the manuscript.

## Supplementary Table, Figures and Datasets

Table S1: Selected circadian clock and clock-associated genes in sorghum

Table S2: Enriched Gene Ontology terms within the module Macia-H14.

Figure S1: Temperature stress responses of the CBF subfamily in sorghum and maize.

Figure S2: Proportions of transcription factors within lists of DEGs.

Figure S3: Temperature stress responses of the HSF family in sorghum and maize.

Figure S4: Proportions of genes with rhythmic expression within temperature stress responsive genes

Figure S5: Enriched phases in lists of DEGs presented in Figure 1B, in thermo-tolerant genotypes SC224 and Macia

Figure S6: Transcript abundance profiles of circadian clock genes.

Figure S7: Summary of the data mining procedure.

Figure S8: Gene expression profiles of modules identified in the co-expression network analysis performed from RTx430 data in the cold stress experiment.

Figure S9: Gene expression profiles of modules identified in the co-expression network analysis performed from SC224 data in the cold stress experiment.

Figure S10: Gene expression profiles of modules identified in the co-expression network analysis performed from RTx430 data in the heat stress experiment.

Figure S11: Gene expression profiles of modules identified in the co-expression network analysis performed from Macia data in the heat stress experiment.

Figure S12: Number of transcription factor (TF) family members within selected network modules.

Supplementary Datasets S1: Normalized (rlog transformation) expression data of sorghum genes

Supplementary Datasets S2: Pairwise comparison analysis Supplementary Datasets S3: Lists of differentially expressed Supplementary Datasets S4: Phase enrichment analysis Supplementary Datasets S5: Likelihood Ratio Test analysis

Supplementary Datasets S6: List of genes with significant interaction between temperature and genotype

Supplementary Datasets S7: Co-expression network modules

Supplementary Datasets S8: List of sorghum EF-hand Ca2+-binding protein family

