## Supplementary Material for "Time of day and genotype sensitivity adjust molecular responses to temperature stress in sorghum"

**Supplementary Table S1. Selected circadian clock and clock-associated genes in sorghum, from Lai et al. (2020).**

| ID | Code |
| --- | --- |
| Sobic.001G411400 | PRR73-LIKE |
| Sobic.002G275100 | PRR95-LIKE |
| Sobic.003G040900 | GI-LIKE |
| Sobic.003G191700 | ELF3-LIKE1 |
| Sobic.004G216700 | TOC1-LIKE |
| Sobic.005G145300 | FKF1-LIKE |
| Sobic.007G047400 | LHY-LIKE |
| Sobic.009G257300 | ELF3-LIKE2 |

**Supplementary Table S2. Enriched Gene Ontology terms within the module Macia-H14**. The online tool 'agriGO v 2.0 was used for this analysis (Tian *et al.* 2017). F, Molecular function; C, Cellular component.

| **GO term** | **Ontology** | **Description** | **Number in input list** | **Number in the Reference** | **P-value** | **FDR** |
| --- | --- | --- | --- | --- | --- | --- |
| GO:0051082 | F | unfolded protein binding | 16 | 33 | 1,00E-05 | 0.01 |
| GO:0051087 | F | chaperone binding | 11 | 18 | 5.2e-05 | 0.026 |
| GO:0043231 | C | intracellular membrane-bounded organelle | 154 | 936 | 1.8e-06 | 0.00036 |
| GO:0043227 | C | membrane-bounded organelle | 154 | 936 | 1.8e-06 | 0.00036 |
| GO:0044428 | C | nuclear part | 36 | 149 | 5,00E-05 | 0.0059 |
| GO:0005654 | C | nucleoplasm | 19 | 56 | 8.6e-05 | 0.0059 |
| GO:0005634 | C | nucleus | 109 | 669 | 8.4e-05 | 0.0059 |
| GO:0044451 | C | nucleoplasm part | 19 | 56 | 8.6e-05 | 0.0059 |
| GO:0070013 | C | intracellular organelle lumen | 24 | 97 | 0.00064 | 0.026 |
| GO:0043233 | C | organelle lumen | 24 | 97 | 0.00064 | 0.026 |
| GO:0031974 | C | membrane-enclosed lumen | 24 | 97 | 0.00064 | 0.026 |
| GO:0031981 | C | nuclear lumen | 22 | 85 | 0.00063 | 0.026 |

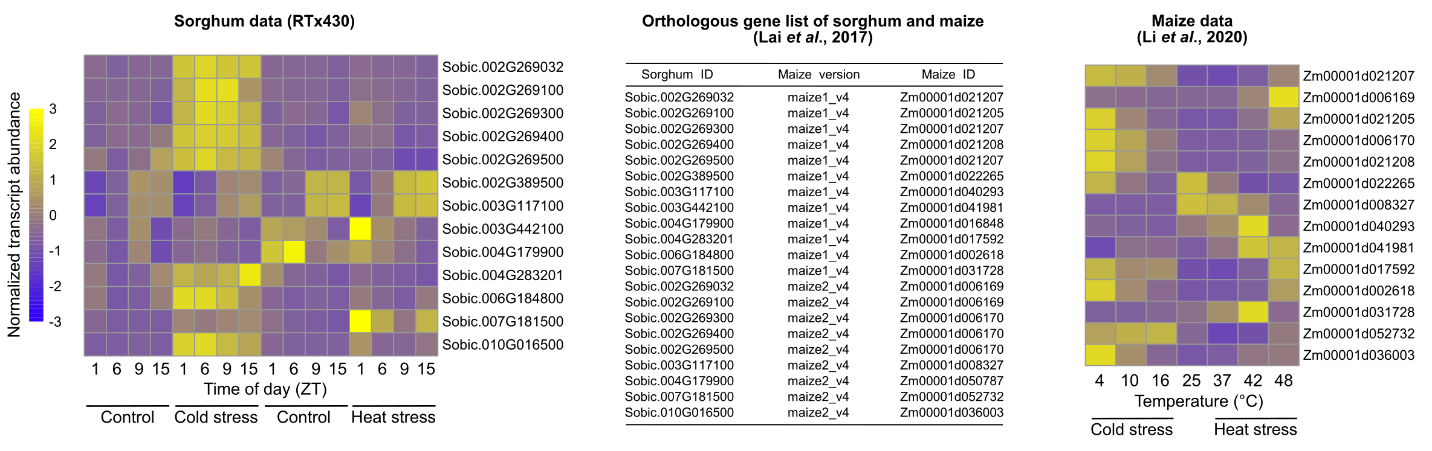

**Supplementary Figure S1. Temperature stress responses of the CBF subfamily in sorghum and maize.** Sorghum data correspond to rlog values in the RTx430 genotype. Orthologous genes between sorghum and maize were downloaded from (Xianjun Lai, Yan & Schnable 2017). Maize data were downloaded from Li et al. (Li *et al.* 2020) and correspond to FPKM values. Data were scaled by row.

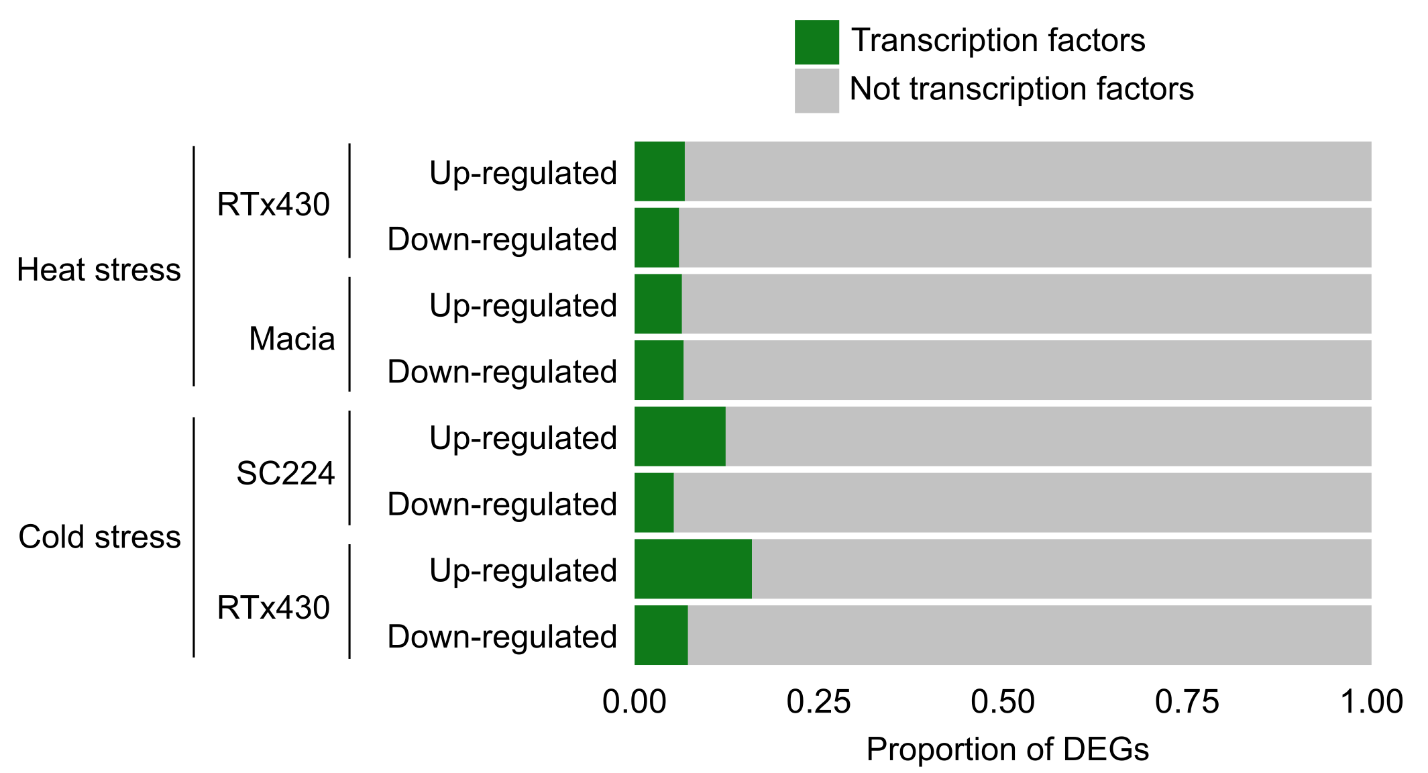

**Supplementary Figure S2. Proportions of transcription factors within lists of DEGs.** Up-regulated and down-regulated mean that the genes were significantly (FDR < 0.05 and Log_2_ Fold Change > |1|) more and less expressed under temperature stress as compared to control conditions, respectively.

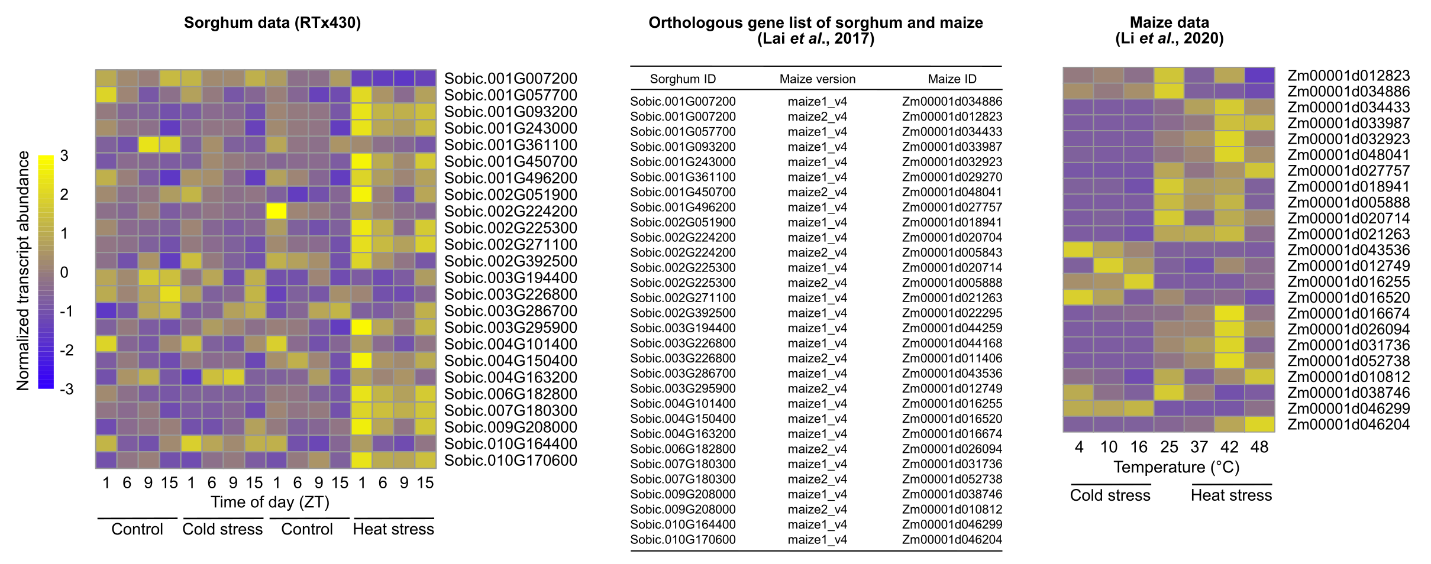

**Supplementary Figure S3. Temperature stress responses of the HSF family in sorghum and maize.** Sorghum data correspond to rlog values in the RTx430 genotype. Orthologous genes between sorghum and maize were downloaded from Xianjun Lai, Yan & Schnable (2017). Maize data were downloaded from Li et al. (2020) and correspond to FPKM values. Data were scaled by row.

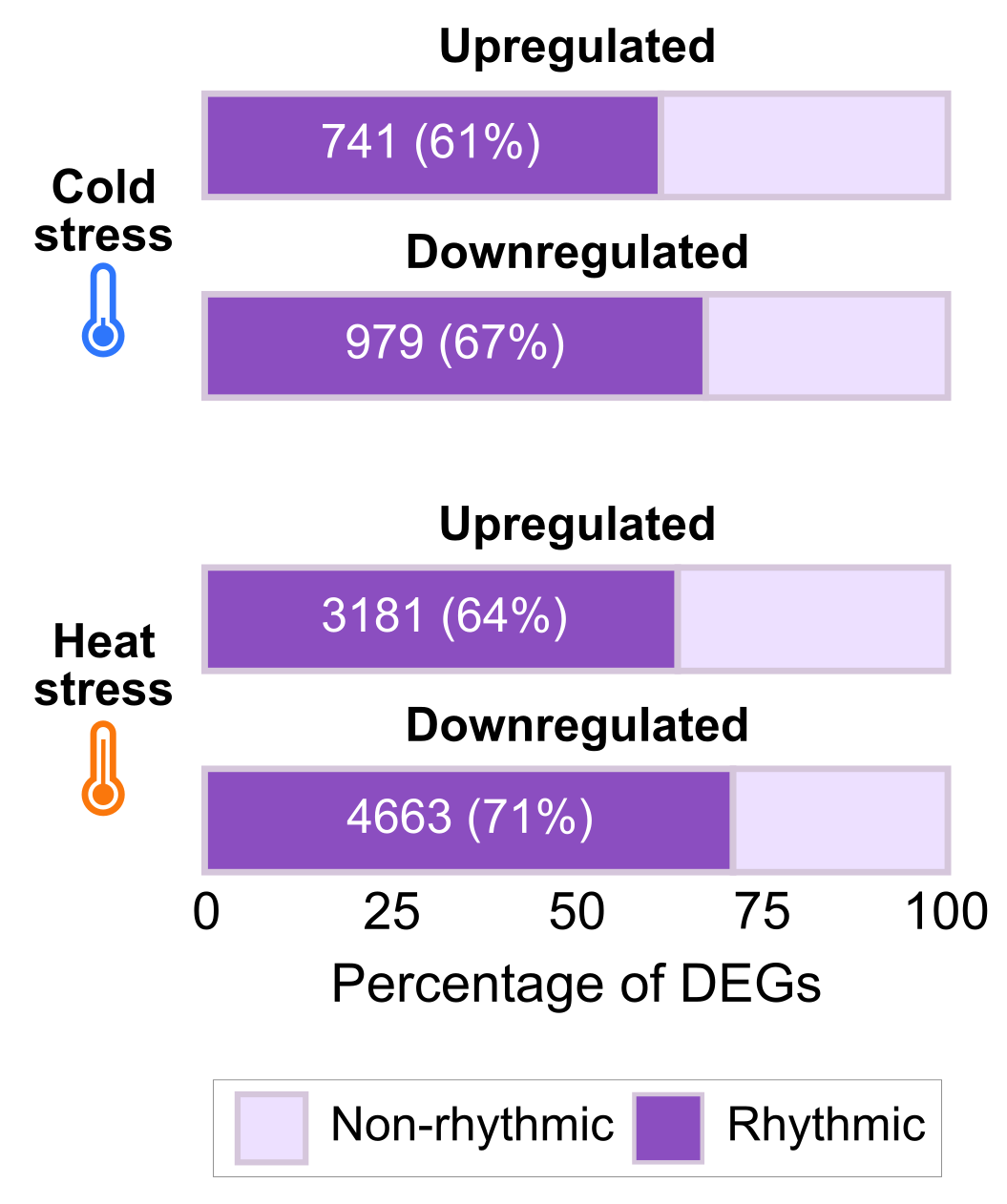

**Supplementary Figure S4. Proportions of genes with rhythmic expression within temperature stress responsive genes.** The list of genes with rhythmic expression was identified in Lai et al. (2020). Upregulated (FDR < 0.05, Log_2_ Fold Change > 1) and downregulated (FDR < 0.05 and Log_2_ Fold Change <(-1)) DEGs were identified by comparing temperature stress vs control conditions, using pairwise comparisons. DEGs correspond to genes with a differential expression either in the thermo-tolerant or susceptible genotype, or in both (presented in Figure 1B).

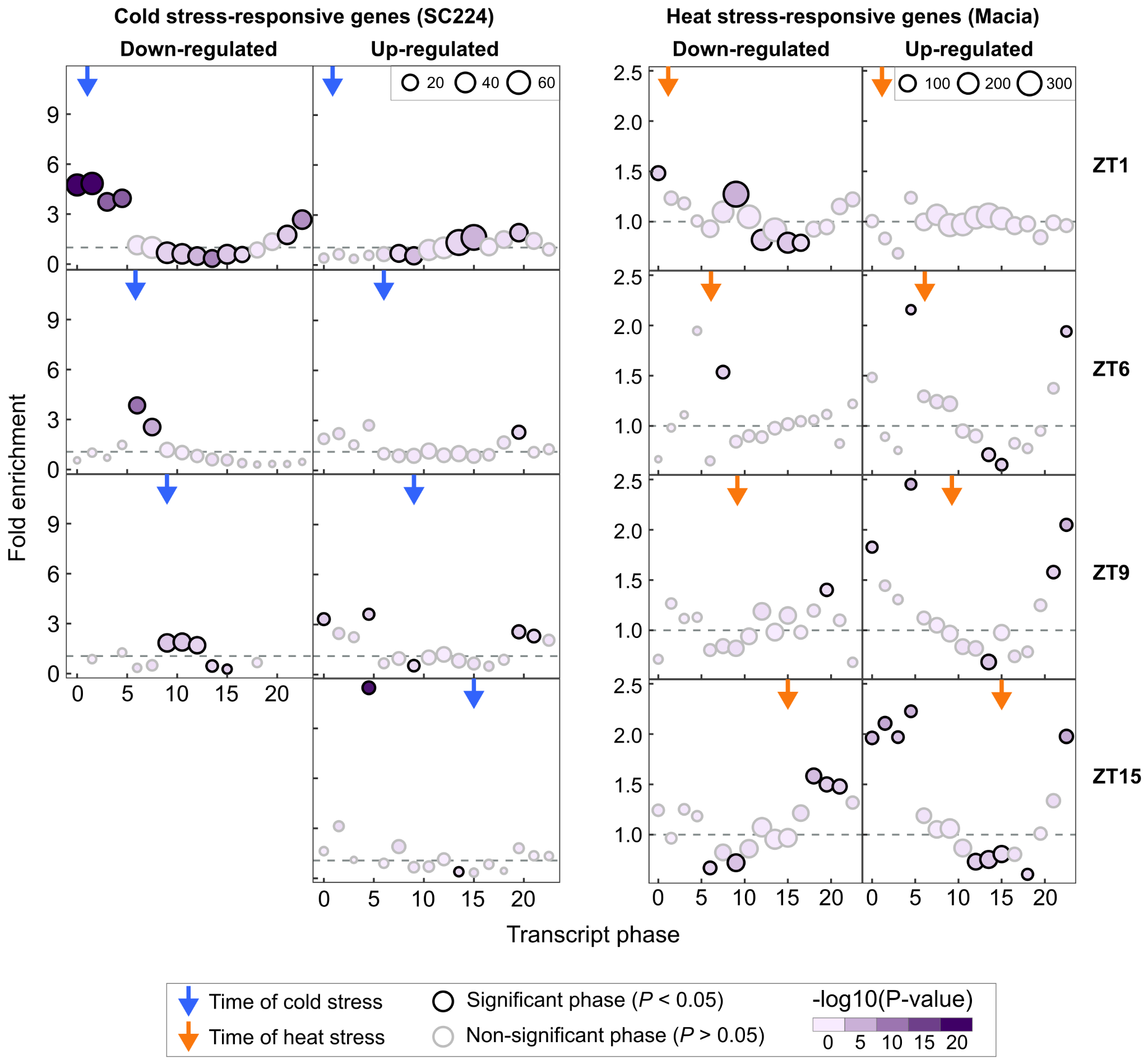

**Supplementary Figure S5. Enriched phases in lists of DEGs presented in Figure 1B, in thermo-tolerant genotypes SC224 and Macia.** The phase is defined as the timing of peak abundance (a phase of 0 and 12 indicates a peak abundance at subjective dawn and the beginning of the subjective night, respectively). Proportions of the different phases in the lists of DEGs were compared to those of all rhythmic genes identified in Lai et al. (2020). Only genes identified as rhythmic in Lai et al. (Lai *et al.* 2020) were considered for this analysis. Horizontal gray dashed lines correspond to a fold enrichment of 1. Bubble plots represent over- (fold enrichment > 1) and under-represented (fold enrichment < 1) phases in the list of DEGs as compared to the reference. Chi-Square tests were performed and significance was judged at P-value < 0.05. For a meaningful enrichment calculation, only sets of ≥ 100 DEGs were considered for this analysis.

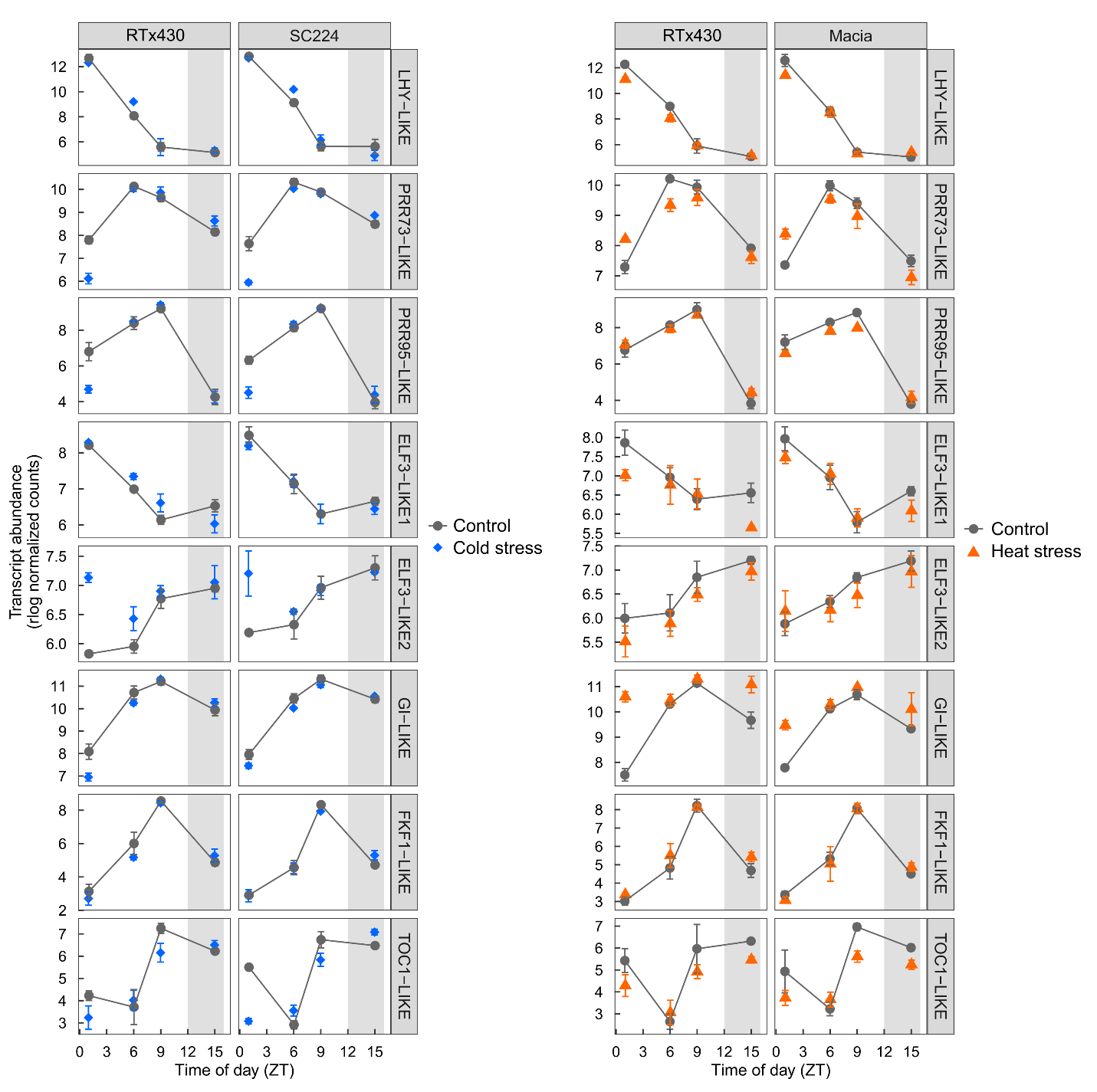

**Supplementary Figure S6. Transcript abundance profiles of circadian clock genes.** Circadian clock genes in sorghum were identified in Lai et al. (2020). Gene IDs are indicated in Supplementary Table S1. Data correspond to rlog (normalized gene expression) values (Means ± SD, *n* = 3). Gray areas represent the night period.

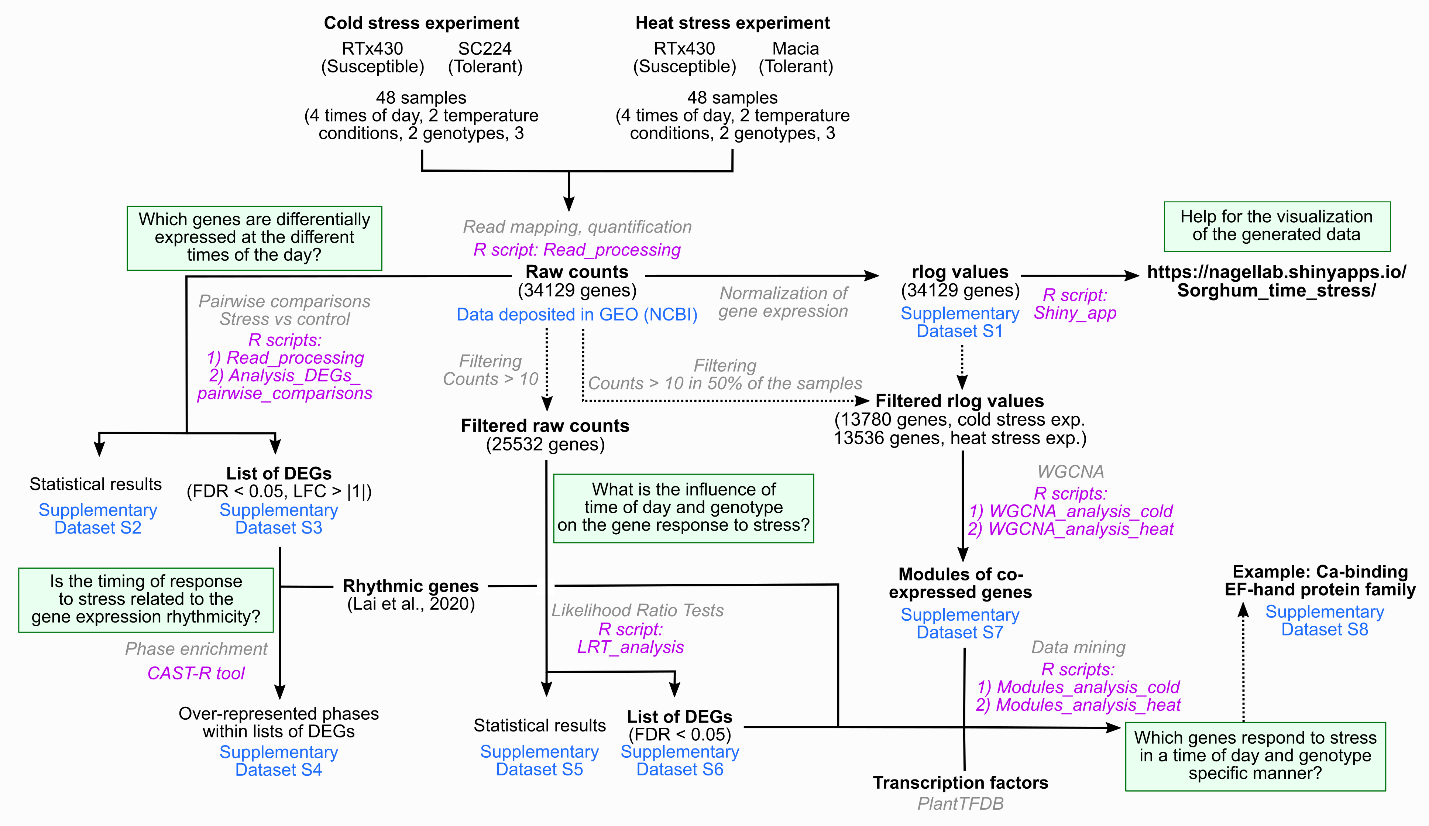

**Supplementary Figure S7. Summary of the data mining procedure.** Questions/objectives are highlighted with gray rectangles. Methods are italicized and written in gray, and corresponding R scripts (available at https://github.com/Nagel-lab/Sorghum_temperature_stress) are italicized and written in purple. Datasets containing results of the different analyses are highlighted in blue. Of note, analyses from the cold stress and heat stress experiments were performed separately.

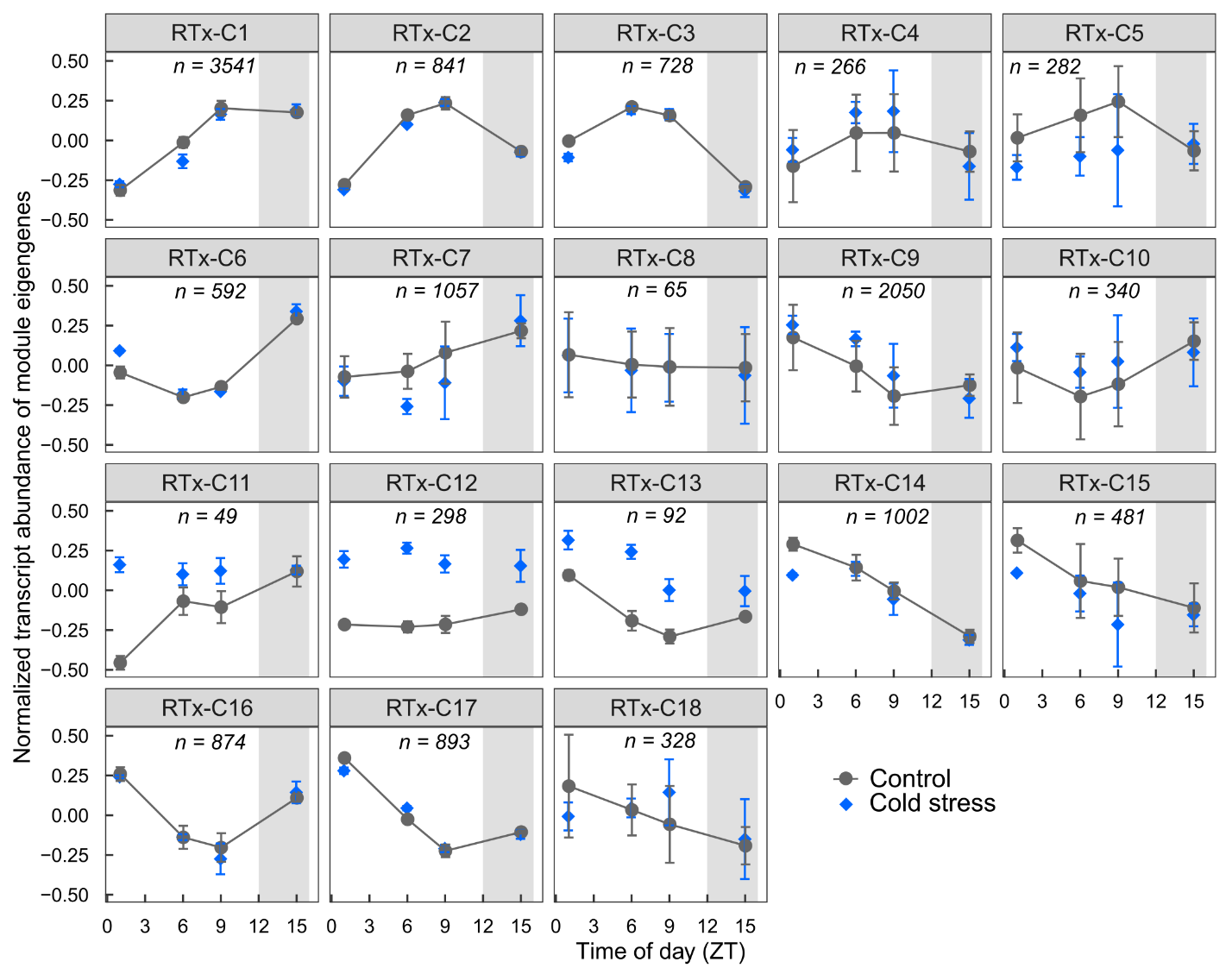

**Supplementary Figure S8. Gene expression profiles of modules identified in the co-expression network analysis performed from RTx430 data in the cold stress experiment.** Profiles were generated from module eigengene data (Means ± SD, *n* = 3). The number of genes within each module is indicated (*e.g. n* = 3541 genes in RTx-C1). Gray areas represent the night period.

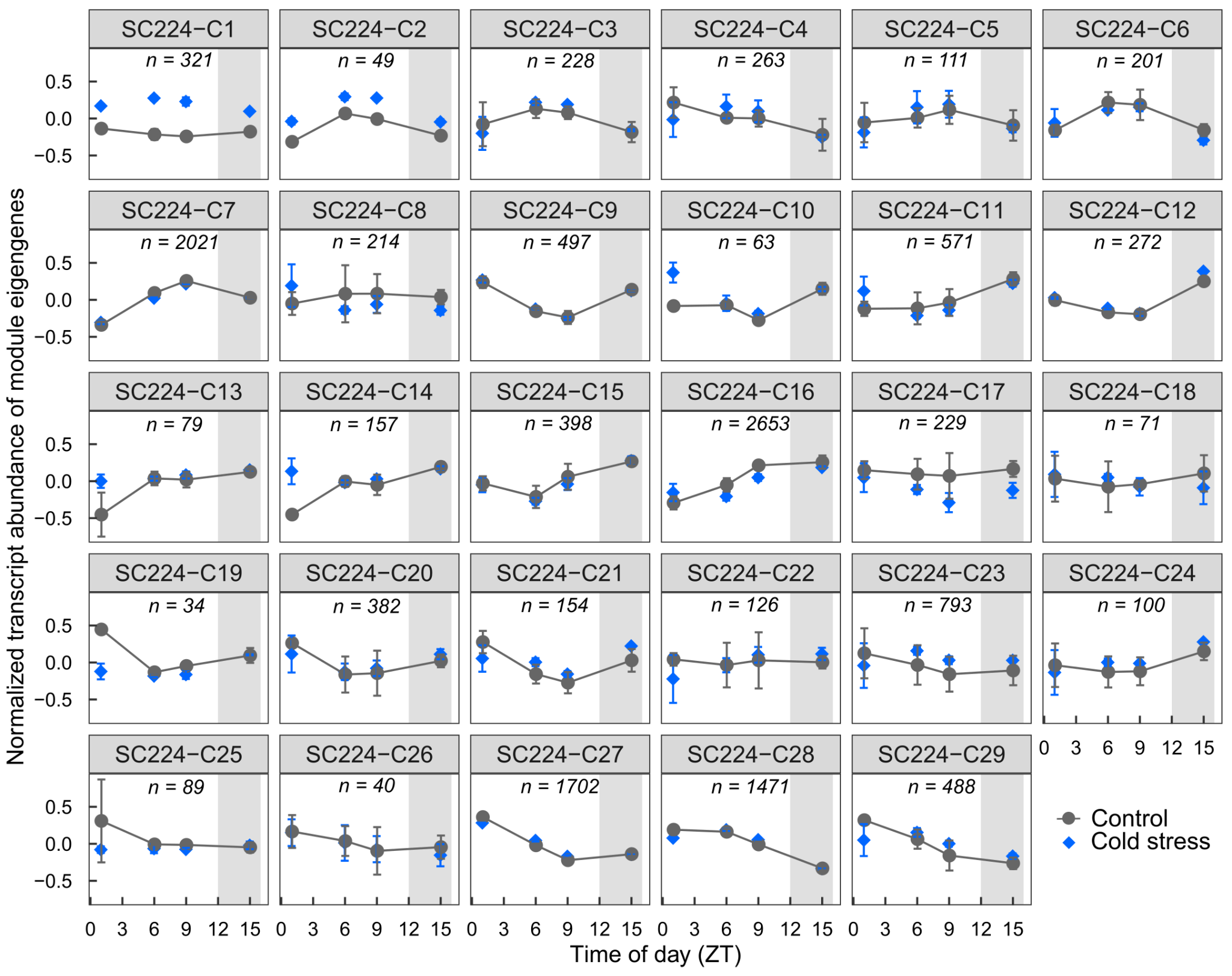

**Supplementary Figure S9. Gene expression profiles of modules identified in the co-expression network analysis performed from SC224 data in the cold stress experiment.** Profiles were generated from module eigengene data (Means ± SD, *n* = 3). The number of genes within each module is indicated (*e.g. n* = 321 genes in SC224-C1). Gray areas represent the night period.

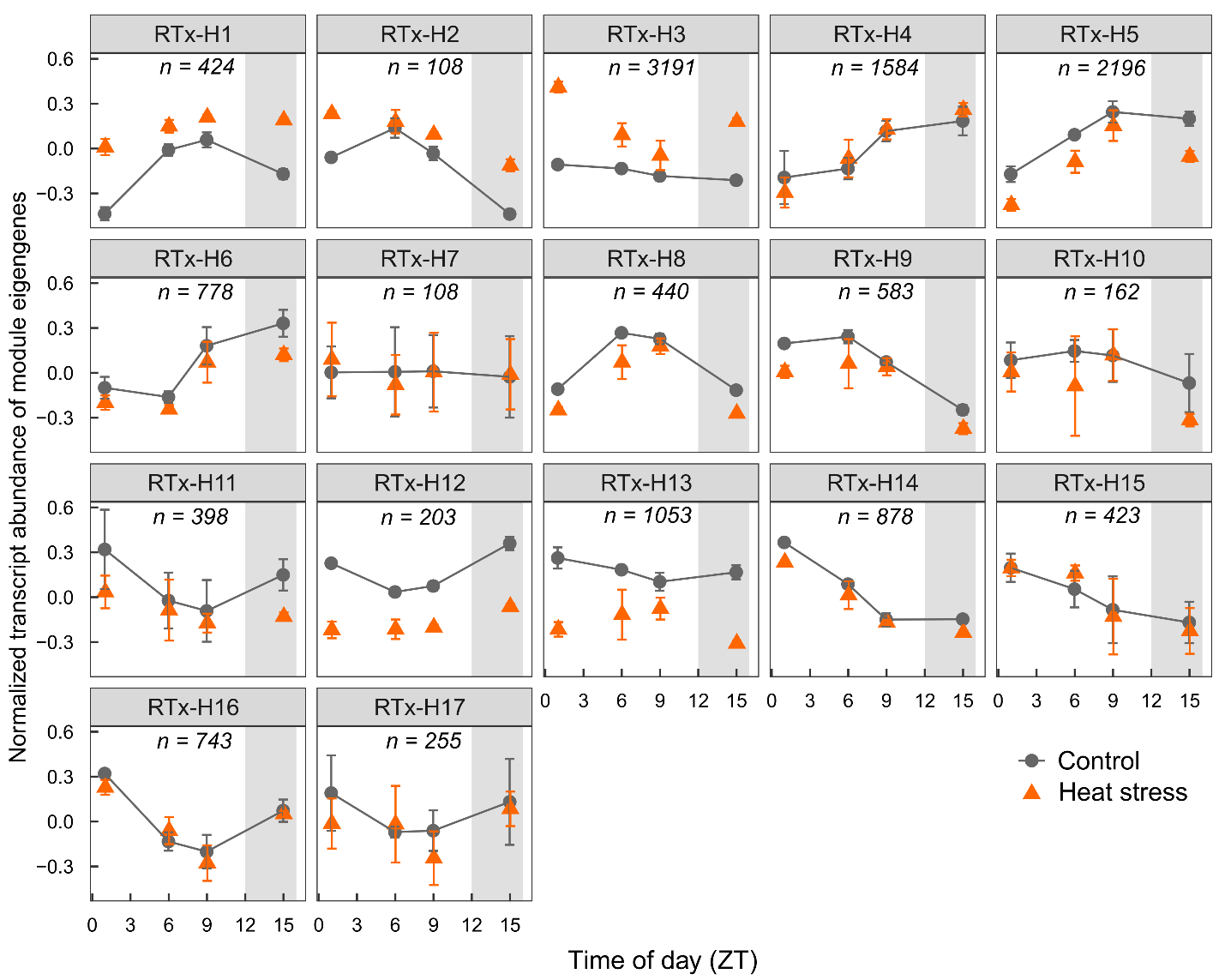

**Supplementary Figure S10. Gene expression profiles of modules identified in the co-expression network analysis performed from RTx430 data in the heat stress experiment.** Profiles were generated from module eigengene data (Means ± SD, *n* = 3). The number of genes within each module is indicated (*e.g. n* = 424 genes in RTx-H1). Gray areas represent the night period.

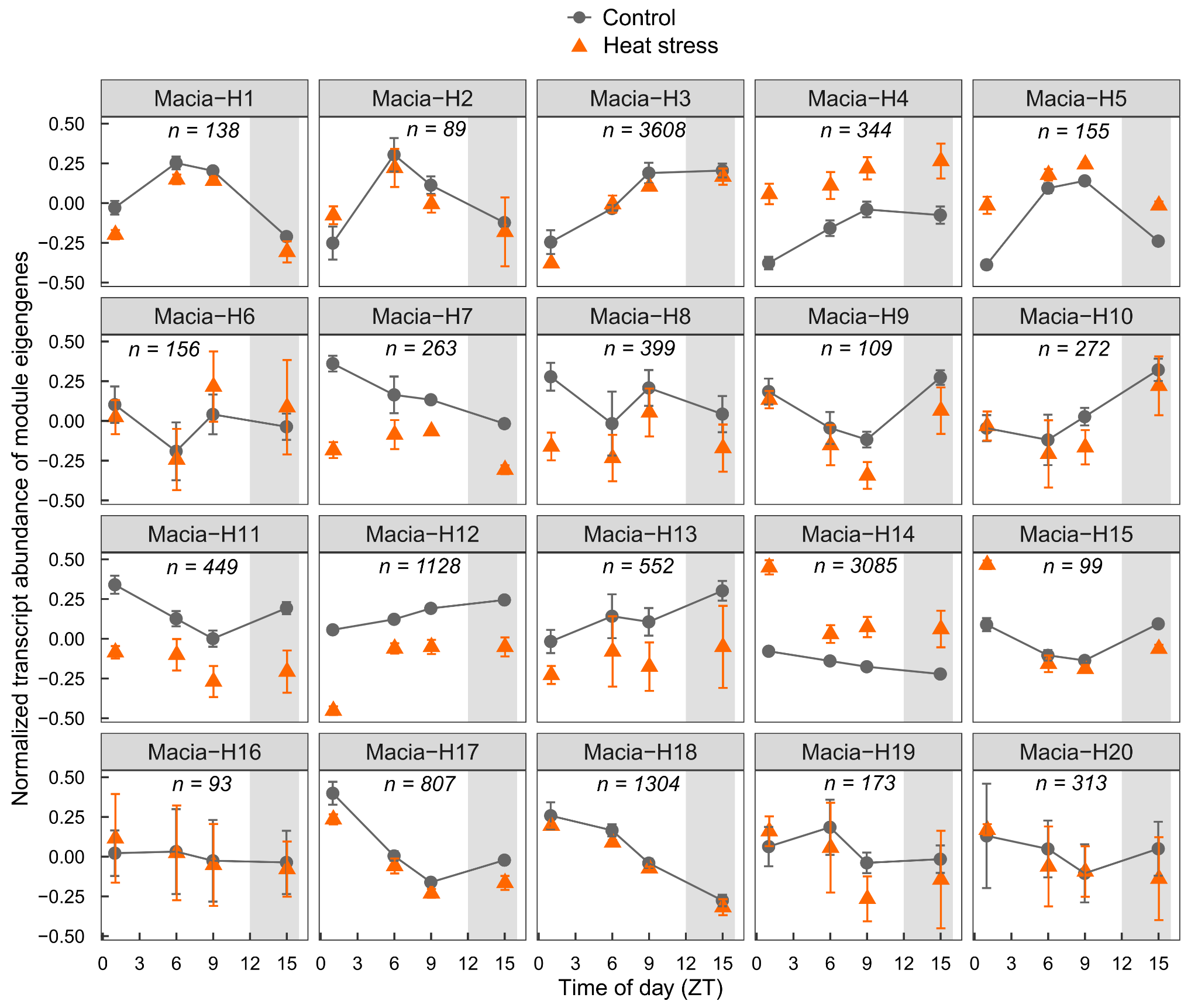

**Supplementary Figure S11. Gene expression profiles of modules identified in the co-expression network analysis performed from Macia data in the heat stress experiment.** Profiles were generated from module eigengene data (Means ± SD, *n* = 3). The number of genes within each module is indicated (*e.g. n* = 138 genes in Macia-H1). Gray areas represent the night period.

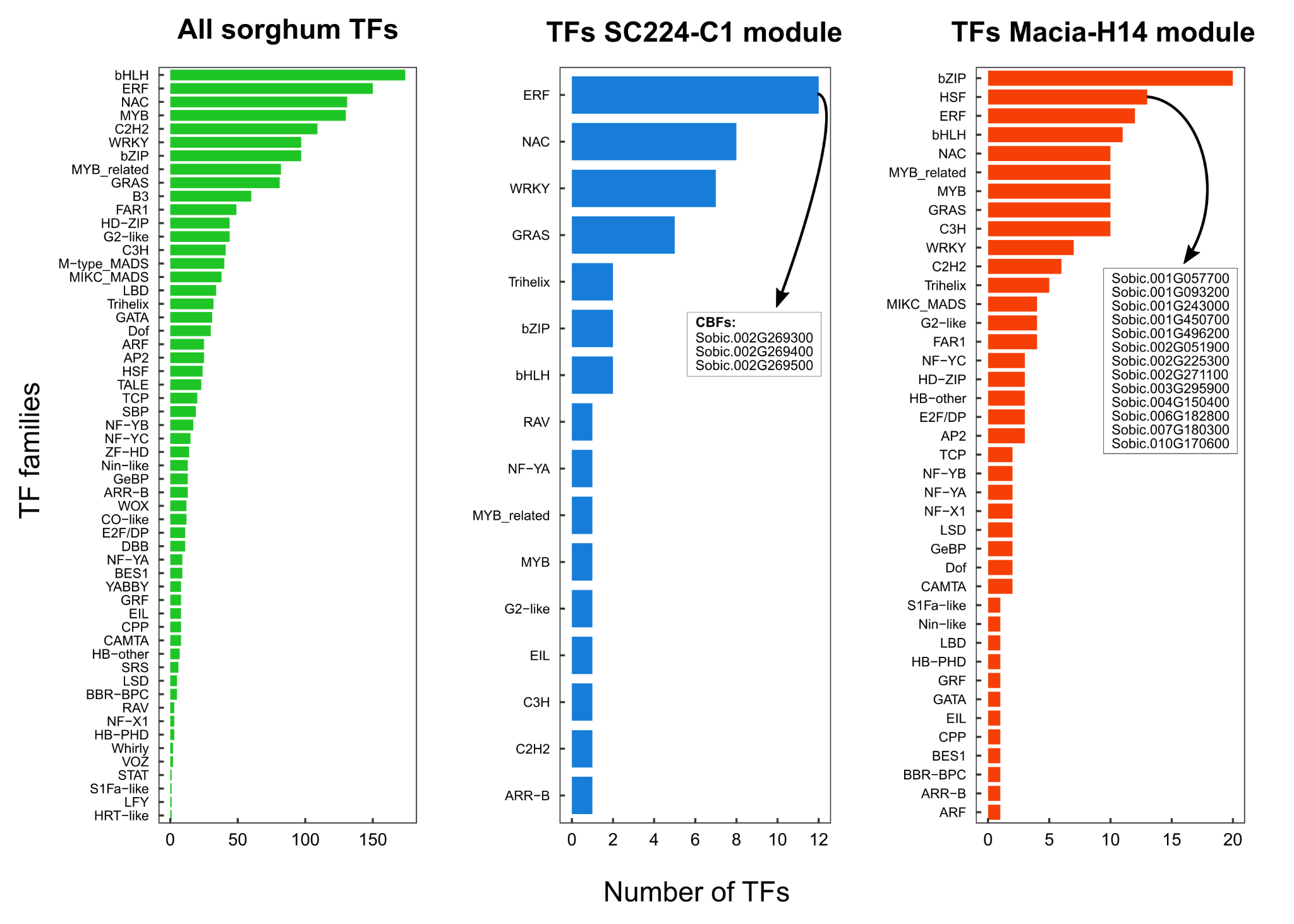

**Supplementary Figure S12. Number of transcription factor (TF) family members within selected network modules.** All sorghum TFs were downloaded from PlantTFDB v 5.0 (Jin *et al.* 2017). Modules SC224-C1 and Macia-H14 are presented in Supplementary Figures S7 and S9, respectively, and highlighted in Figure 4. Gene IDs for CBFs and HSFs genes are indicated in SC224-C1 and Macia-H14, respectively. CBFs are a subfamily of the ERF TF family.
